# Extreme genome diversity and cryptic speciation in a harmful algal bloom forming eukaryote

**DOI:** 10.1101/2022.10.25.513657

**Authors:** Jennifer H. Wisecaver, Robert P. Auber, Amanda L. Pendleton, Nathan F. Watervoort, Timothy R. Fallon, Olivia L. Riedling, Schonna R. Manning, Bradley S. Moore, William W. Driscoll

**Author notes:** These authors contributed equally (Robert P. Auber, Amanda L. Pendleton). Correspondence: Jennifer Wisecaver, 175 S University St., West Lafayette, IN 47907; e.

## Abstract

Harmful algal blooms (HABs) of the toxic haptophyte *Prymnesium parvum* are a recurrent problem in many inland and estuarine waters around the world. Strains of *P. parvum* vary in the toxins they produce and in other physiological traits associated with HABs, but the genetic basis for this variation is unknown. To investigate genome diversity in this morphospecies, we generated genome assemblies for fifteen phylogenetically and geographically diverse strains of *P. parvum* including Hi-C guided, near-chromosome level assemblies for two strains. Comparative analysis revealed considerable DNA content variation between strains, ranging from 115 Mbp to 845 Mbp. Strains included haploids, diploids, and polyploids, but not all differences in DNA content were due to variation in genome copy number. Haploid genome size between strains of different chemotypes differed by as much as 243 Mbp. Syntenic and phylogenetic analyses indicate that UTEX 2797, a common laboratory strain from Texas, is a hybrid that retains two phylogenetically distinct haplotypes. Investigation of gene families variably present across strains identified several functional categories associated with metabolism, including candidates for the biosynthesis of toxic metabolites, as well as genome size variation, including recent proliferations of transposable elements. Together, our results indicate that *P. parvum* is comprised of multiple cryptic species. These genomes provide a robust phylogenetic and genomic framework for investigations into the eco-physiological consequences of the intra- and inter-specific genetic variation present in *P. parvum* and demonstrate the need for similar resources for other HAB-forming morphospecies.

**SIGNIFICANCE STATEMENT:** Harmful algal blooms (HABs) are a global concern. Efforts to understand the genetic basis of traits associated with the success of HAB-forming species are limited by a dearth of genomic resources. In this paper we present genomes for fifteen strains of *Prymnesium parvum*, a toxic alga that causes ecosystem and societally disruptive HABs around the world. We uncover an unprecedented amount of sequence-level, gene family, and genome architecture evolution in *P. parvum* and provide evidence for both cryptic speciation and hybridization. These results illustrate how both inter- and intraspecific genetic variation can be dramatically underestimated in a protist morphospecies. More work is needed to understand the eco-physiological consequences of hidden genetic diversity in *P. parvum* and HAB-forming species more generally.

## INTRODUCTION

Environmentally disruptive harmful algal blooms (HABs) are increasing in frequency and severity in many marine, brackish, and freshwater ecosystems (1–3). Knowledge of genetic variation within HAB populations can aid efforts to predict when HABs will occur and model HAB impacts in a changing climate. However, genome-level data are currently limited for HAB-forming species (4). One such species, *Prymnesium parvum*, is a unicellular microalga belonging to the haptophyte clade of eukaryotes (5). Globally, HABs of *P. parvum* have considerable ecologic and economic impacts. In the southwestern United States, particularly in Texas, blooms are correlated with several troubling environmental trends including increased use of glyphosate herbicides, increased concentrations of atmospheric CO_2_, and decreased wetland cover (6, 7).

*P. parvum’s* success as a HAB-forming species likely results from several interrelated aspects of its biology. The species is extremely eurythermal and euryhaline, tolerating temperatures ranging from 2 to 32 °C and salinities ranging from 0.5 to 125% of typical seawater (8). Although *P. parvum* is photosynthetic and maintains permanent chloroplasts, it is also a mixotroph and able to ingest organic material via phagocytosis (9, 10). Despite its small size, *P. parvum* is a ferocious facultative predator capable of swarming and killing various prey including bacteria, other microbial eukaryotes, aquatic invertebrates, and even fish (11–14). The species produces toxins that are thought to assist in this predatory behavior and/or inhibit the activity of competitors and grazers (5). Heterotrophy and toxicity can increase in *P. parvum* cultures grown in sub-optimal conditions, *e.g*., when nutrients are limiting or water temperature/salinity is at the edge of what a strain can tolerate (8, 10, 15).

*P. parvum* produces a variety of toxic metabolites (16). Most conspicuous are the prymnesins, massive ladder-frame polyether compounds that are unique to *P. parvum* but structurally similar to the phycotoxins of many HAB-forming dinoflagellates (17). More than 50 different prymnesins have been identified and are grouped into types based on the number of carbons in their aglycone backbone. The backbone of A-type prymnesins contain 91 carbons, while B-type and C-type prymnesins contain 85 and 83 carbons, respectively (18, 19). To date, all strains of *P. parvum* produce just one type of prymnesin, and strains that produce the same prymnesin type (*i.e*., chemotype) group together in phylogenies of rDNA sequences (19, 20). The strong agreement between phylogeny and chemotype—despite strains of different chemotypes often existing in the same geographical location—suggests that there is limited gene flow between chemotypes of *P. parvum* (19).

Strains of *P. parvum* may appear morphologically indistinguishable, but the species harbors considerable phenotypic diversity. In addition to the previously discussed diversity in prymnesin chemotypes, the amount of prymnesin produced varies between strains (20) as does overall toxicity (20–23). Predatory behavior also varies between *P. parvum* strains, even between strains taken from the same bloom (24). Moreover, several abiotic factors including temperature, salinity, pH, and irradiance have distinct effects on the relative growth rate and toxicity of different strains (23, 25–27), demonstrating remarkable genotype by environment interactions and adaptive plasticity in *P. parvum*.

Extensive physiological and biochemical variation between strains of *P. parvum* indicates either the existence of multiple cryptic species (19) or extreme standing genetic variation in this morphospecies. However, the genetic differences between strains have not yet been quantified. Here, we generated fifteen sequenced genomes of *P. parvum* strains and report considerable genomic variation, both at the nucleotide level and in terms of gene family presence/absence. Our comparative phylogenetic analysis supports monophyletic origins of A-, B-, and C-type prymnesins, with C-type strains having more within-clade sequence diversity compared to A- and B-types. Moreover, we found that the three prymnesin chemotypes of *P. parvum* have dramatically different haploid genome sizes. In addition to haploid strains identified in all three chemotypes, we also identified A- and C-type diploid strains, as well as A- and B-type 4n tetraploids. Lastly, we present evidence that UTEX 2797, a common laboratory strain from Texas, is a hybrid that retains two phylogenetically distinct haplotypes.

## RESULTS

### *Prymnesium parvum* genome assemblies

We obtained Hi-C scaffolded, highly contiguous genome assemblies of two *P. parvum* strains from Texas: UTEX 2797 and 12B1. UTEX 2797 was selected for sequencing due to its use in numerous toxicology and physiology experiments of *P. parvum* (*e.g*., 19, 20, 22, 27–32). However, preliminary analysis of k-mer frequencies revealed that UTEX 2797 displays high sequence-level heterozygosity (Fig. 1A), which can complicate genome assembly. Therefore, we also selected strain 12B1 (21), which had no observable heterozygosity to obtain a homozygous reference assembly (Fig. 1A). UTEX 2797 produces A-type prymnesins (18, 19), and our chemotyping analysis revealed strain 12B1 produces A-type prymnesins as well (Fig. S1).

**Figure 1.**
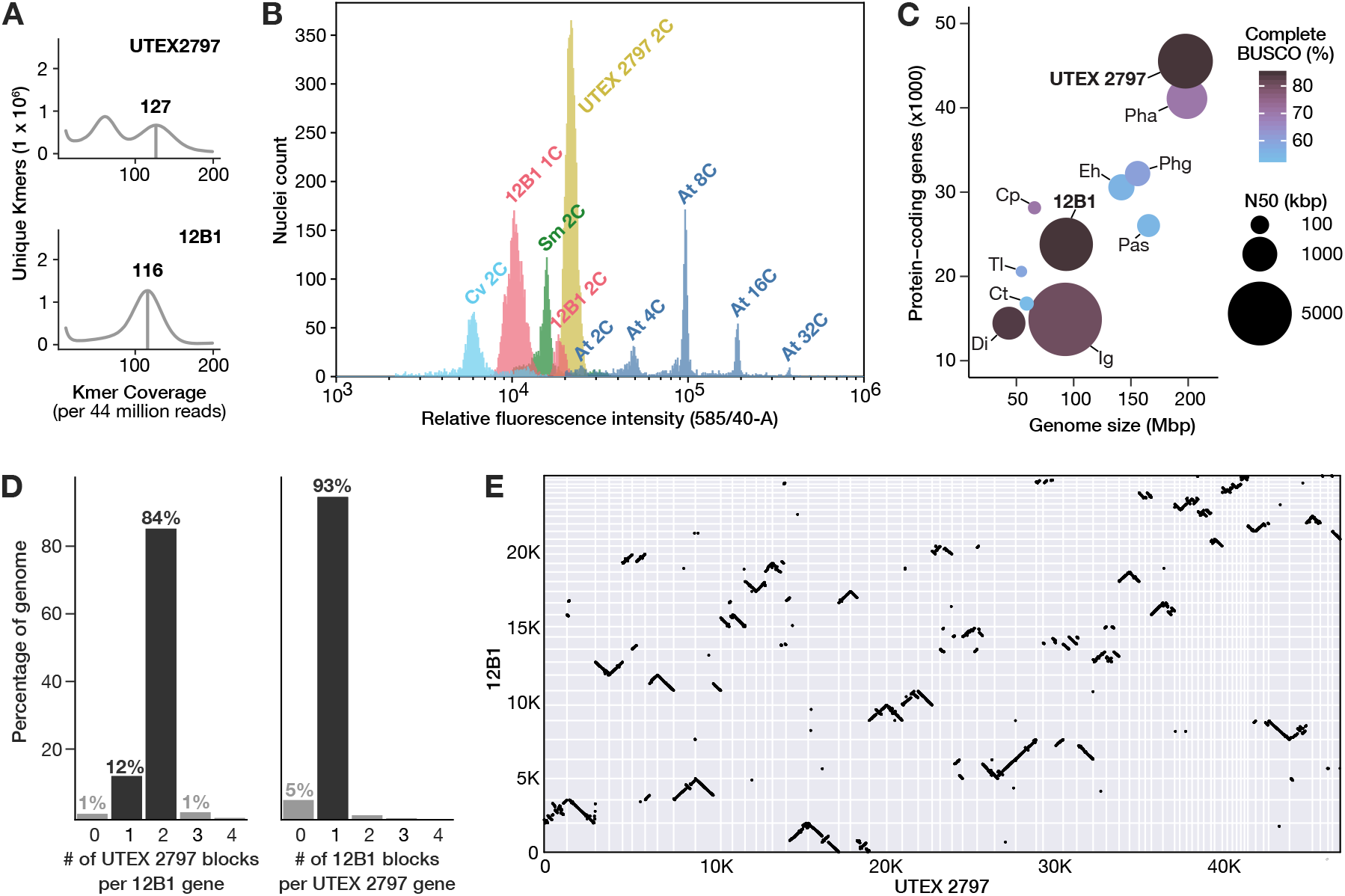
Genome metrics for *P. parvum* Hi-C scaffolded long-read assemblies. A) K-mer frequency plots showing estimated heterozygosity for strains UTEX 2797 and 12B1.The homozygous k-mer peaks are indicated by vertical bars, with the number above the homozygous peaks indicating the coverage of maximal unique k-mers (CMUKs). UTEX 2797 has high heterozygosity as indicated by the large second peak at half the k-mer coverage of the homozygous peak. B) Histogram indicating the relative fluorescent intensity of propidium iodide stained nuclei for strains 12B1 (pink) and UTEX 2797 (yellow) relative to genome standards: Cv, *Chlorella vulgaris* (light blue); Sm, *Selaginella moellendorfii* (green); At, *Arabidopsis thaliana* Col-0 (dark blue). The endoreduplicative nature of the *A. thaliana* tissue enabled identification of 2C, 4C, 8C, 16C, and 32C nuclei in this species (62). C) Comparison of assembly completeness and contiguity across eleven haptophyte genome assemblies. Cp, *Chrysochromulina parva*; Ct, *Chrysochromulina tobinii*; Dl, *Diacronema lutheri*; Eh, *Emiliania huxleyi*; Ig, *Isochrysis galbana*; Pas, Pavlovales sp.; Pha, *Phaeocystis antarctica*; Phg, *Phaeocystis globosa*; Tl, *Tisochrysis lutea*. D) Ratio of syntenic block counts between *P. parvum* genome assemblies for strains 12B1 and UTEX 2797. Syntenic blocks of UTEX 2797 per 12B1 gene (left) and syntenic blocks of 12B1 per UTEX 2797 gene (right) are shown, indicating a clear 1:2 pattern of 12B1 to UTEX 2797. E) Macrosynteny of the 12B1 and UTEX 2797 genomes. Syntenic gene pairs are denoted by black points and positionally oriented by scaffold (grid squares).

The nuclear DNA content of both strains was estimated using flow cytometry (Fig. 1B). The majority of 12B1 nuclei contained an average of 0.12 pg DNA, which corresponds to approximately 115.6 million base pairs (Mbp), assuming a conversion factor of 1 pg = 0.98 × 10^9^ base pairs (33). The 12B1 flow cytometry histogram also contained a second, substantially smaller peak corresponding to approximately 0.23 pg DNA (228 Mbp), indicating that some cells may have been in the process of dividing (*i.e*., the G2 phase of the cell cycle) (Fig. 1B). The UTEX 2797 flow cytometry histogram contained a single dominant peak corresponding to 0.28 pg DNA, or 274.4 Mbp (Fig. 1B, Table S1). A cryptic sexual lifecycle for *P. parvum* has been proposed that alternates between haploid and diploid forms, which are only distinguishable by transmission electron microscopy of the cells’ outer scales (34). However, neither syngamy nor meiosis have been observed in *P. parvum*, and its lifecycle remains unknown. Therefore, we must be cautious when inferring the haploid genome size and relative ploidy states of different strains, and we tentatively labeled the dominant peaks in 12B1 and UTEX 2797 as 1C and 2C, respectively (Fig. 1B).

The Hi-C scaffolded genome assembly for 12B1 consisted of 34 scaffolds spanning 93.6 Mbp with an N50 of 3.2 Mbp (Table S2). The UTEX 2797 assembly was over twice the length of the 12B1 assembly at 197.6 Mbp and consisted of 66 scaffolds with an N50 of 3.4 Mbp (Table S2). Compared to nine other haptophyte assemblies (see Table S3 and references therein), the UTEX 2797 and 12B1 assemblies are the second and third most contiguous (Fig. 1C). Chromosomal end-to-end assembly was accomplished for 29.4% and 7.5% of 12B1 and UTEX 2797 scaffolds, respectively (Table S4), evidenced by the presence of TTAGGG telomeric sequences observed in other haptophytes (35). Overall, most 12B1 scaffolds (73.5%) had at least one predicted telomere. In contrast, only 42.4% of UTEX 2797 scaffolds had any predicted telomeres (Fig. S2), which suggests that the level of heterozygosity in the UTEX 2797 genome may have impacted the complete assembly of these repetitive regions. Total repetitive sequence makes up 29.4% and 35.5% of the 12B1 and UTEX 2797 genome assemblies, respectively (Table S5), which are within the range of values (22.9% - 64%) reported for other haptophytes (36–38).

We predicted and annotated 23,802 and 45,535 protein-coding genes in the genomes of 12B1 and UTEX 2797, respectively (Table S6). In total, 86.2% and 86.4% of the 12B1 and UTEX 2797 genes could be assigned predicted functional annotations through InterProScan (39). The completeness of the predicted gene sets were assessed with BUSCO (40); 216 of 255 conserved eukaryotic genes (84.7%) were complete within both predicted proteomes (Table S7). This level of BUSCO recovery is the greatest of any currently available haptophyte genome assembly (Fig. 1C, Table S3). While only 5.1% of BUSCOs were duplicated in 12B1, the majority of BUSCOs (75.5%) were duplicated in UTEX 2797 (Table S7).

Synteny and collinearity analyses further highlight the duplicated state of the UTEX 2797 genome assembly. A strong 2:1 synteny pattern is observed between the genes of UTEX 2797 and 12B1; 93% of UTEX 2797 genes are syntenic to a single region in 12B1, whereas 84% of 12B1 genes are syntenic to two regions in the UTEX 2797 genome (Fig. 1D). Most 12B1 scaffolds are collinear with two syntenic regions in the UTEX 2797 assembly, but an abundance of large structural variants (*e.g*., inversions, indels, translocations) are apparent both between the two reference strains as well as between syntenic scaffolds of UTEX 2797 (Fig. 1E). Together, these data indicate that the assembly of UTEX 2797 consists of two largely non-collapsed haplotypes, which is consistent with the strain being a highly heterozygous diploid. Due to the extreme heterozygosity present in UTEX 2797, we suspected that the strain may have arisen via hybridization of two divergent parents. However, further investigation required data from additional *P. parvum* strains.

To enable phylogenetic analysis of *P. parvum* and investigate the origin of the divergent haplotypes in UTEX 2797, we selected thirteen additional *P. parvum* strains for genomic characterization that represent different geographical locations and prymnesin chemotypes (Table S8). In addition to 12B1 and UTEX 2797, we included four other strains that produce A-type prymnesins: 12A1, CCMP2941, CCMP3037, and RCC3703 (Fig. S1) (19). Three strains in the analysis produce B-type prymnesins (K-0081, K-0374, and KAC-39), and four produce C-type prymnesins (K-0252, RCC191, RCC1433, and RCC1436) (18, 19). Two additional strains in our analysis (RCC3426 and UTEX 995) have not yet been chemotyped. We assembled and annotated the genomes of these thirteen additional strains using Illumina short reads (Table S2). Genome assembly lengths ranged from 77.0 Mbp (CCMP3707) to 92.9 Mbp (RCC1436). As expected of short-read assemblies, genome contiguity was low, with contig N50s that ranged from 3414 bp (K-0081) to 8872 bp (RCC1433). Nevertheless, coding gene space was well represented, and the number of protein coding genes ranged from 26,458 (CCMP3037) to 32,339 (UTEX 995) (Table S6). BUSCO scores ranged from 65% (UTEX 995) to 81% (RCC1433), comparable to BUSCO scores observed in other Illumina-only haptophyte genome assemblies (Table S7). However, it is notable that 10 unique BUSCOs were not recovered in any of the 24 haptophyte genomes investigated (Table S9). This suggests that some BUSCOs may be absent in haptophytes or too divergent to be recovered using the current BUSCO sequence models.

### High sequence divergence within a unicellular morphospecies

To determine the relatedness among strains of different chemotypes, we first identified orthologous gene families (*i.e*., orthogroups) using OrthoFinder (41). We selected 2699 single-copy orthogroups (SCOGs) present in all 15 strains for concatenation- and coalescent-based species tree analyses. Because the most closely related haptophyte species with sequenced genomes were too divergent to serve as outgroups for a nucleotide-based phylogeny, a nonreversible substitution model was used to evaluate support for different root locations on the *P. parvum* species tree (42). The root with the highest likelihood score caused the C-type prymnesin producers to be paraphyletic, which is a phylogenetic pattern also observed in a prior 18S rDNA phylogeny (20). However, ‘rootstrap’ support for this root location was low (68%), and two alternative root locations, including one in which C-type producers were monophyletic, could not be rejected (AU tree topology test P-values > 0.1; Fig. 2A, Fig. S3).

**Figure 2.**
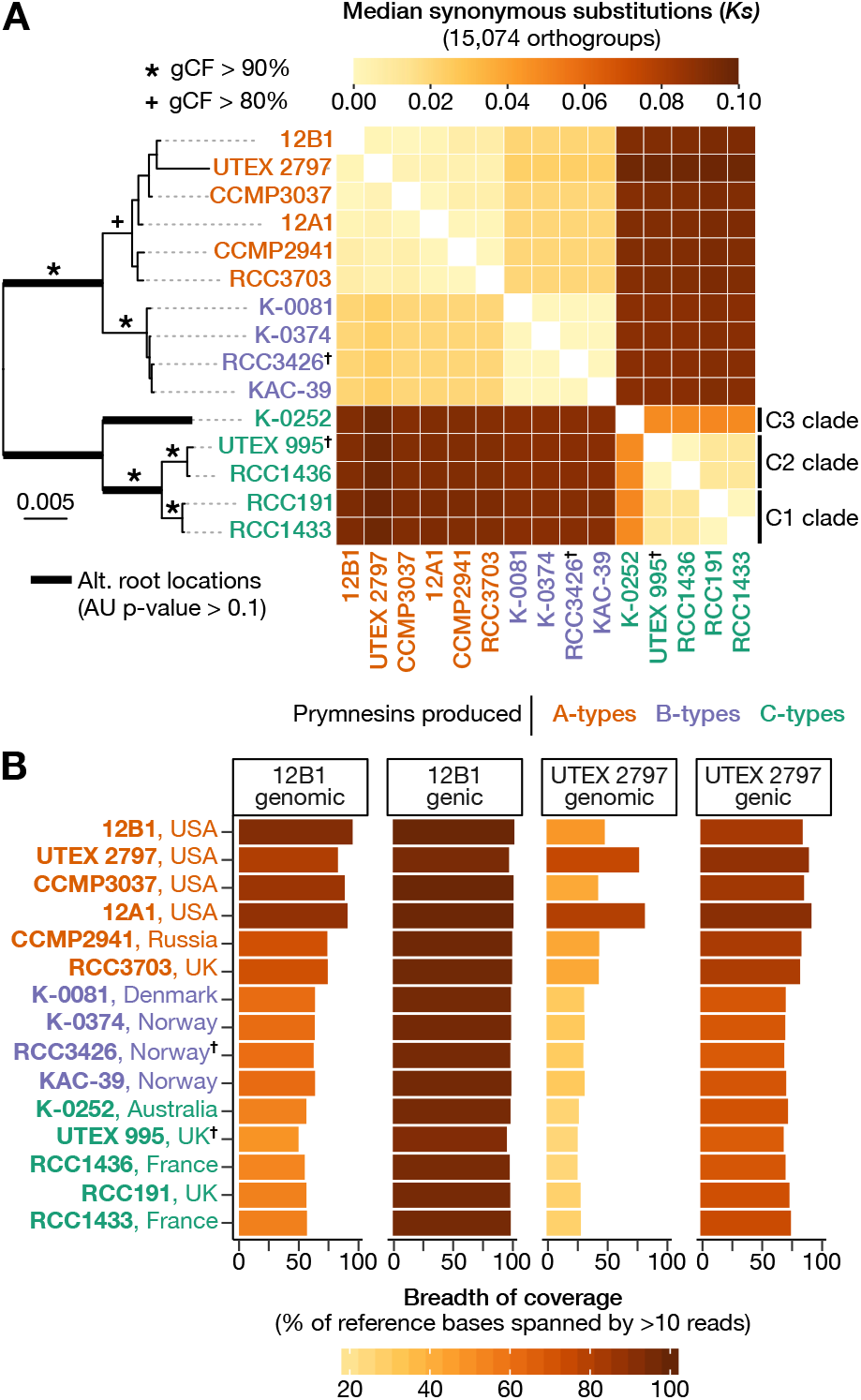
Phylogenomic and breadth of coverage analyses in *P. parvum*. A) Concatenation-based ML species phylogeny showing three possible root locations (bold branches). The heatmap shows the median synonymous substitutions per synonymous site (*Ks*) between all strains. C) Breadth of coverage (BOC) bar plots showing the percent of 12B1 and UTEX 2797 reference bases spanned by >10 Illumina reads from each strain (See Tables S11). BOC is provided for all genomic positions and genic space as separate statistics. The chemotypes of strains RCC3426 and UTEX 995 (†) are inferred based on phylogenetic placement.

Regardless of the root location, both concatenation- and coalescent-based species trees supported the monophyletic origin of A-type and B-type prymnesins (Fig. 2A, Fig. S8). Support for the species tree topology was assessed using gene and site concordance factors (gCF/sCF) (43). Support was generally high across the tree but low within A-type and B-type clades (Fig. 2A, Fig. S8). These low support values also correspond to areas of disagreement between the concatenation and coalescent based phylogenies (Fig. S8). Untyped strain RCC3426 grouped with strains that produce B-type prymnesins with strong support (gCF = 95.1; sCF = 96.7), suggesting RCC3426 also produces B-type prymnesins. Similarly, untyped strain UTEX 995 grouped with C-type strain RCC1436 (gCF = 92.7; sCF = 94.9), indicating it likely produces C-type prymnesins.

Divergence among strains was evaluated using two approaches. First, divergence was estimated based on the average number of substitutions per synonymous site (*Ks*) between gene pairs in 15,074 orthogroups. These orthogroups were selected based on their presence in 10 or more strains and robust nucleotide alignments. Average *Ks* within A- and B-type clades was extremely low at 0.003 and 0.000, respectively (Fig. 2A, Fig. S4, Table S10). The C-type strains formed three distinct groups based on *Ks*: clade C1 (RCC191 and RCC1433), clade C2 (RCC1436, UTEX 995), and clade C3 (K-0252). Median *Ks* was elevated when C-type strains were compared to each other (*Ks* = 0.0241) with the C3 strain, K-0252, from Australia acting as a significant outlier when compared to other C-types (*Ks* = 0.048) (Fig. 2A, Fig. S4, Table S10). The largest *Ks* values occurred when C-type strains were compared to A- and B-types (*Ks* = 0.093). These results suggest that C-type strains form a monophyletic group and that the best root location for the *P. parvum* species tree is along the branch separating C-type strains from A- and B-types (Fig. 2A).

A second approach, complimentary to sequence divergence and based on breadth of coverage (BOC) of each strain’s Illumina reads, was used to explore strain relatedness. Reads were aligned to the 12B1 and UTEX 2797 reference genomes, and BOC was defined as the percentage of reference sequence spanned by greater than ten reads. Average BOC of the haploid 12B1 genome varied dramatically between chemotypes, ranging from 91.7% in A-types, 72.7% in B-types, and 63.4% in C-type strains (Fig. 2B, Table S11). A-type strains from the U.S.A had a higher average BOC (95.4%) compared to the two European strains (84.2%). BOC data from all strains plotted across the 34 scaffolds of the 12B1 reference assembly generally depicted decreased coverage as phylogenetic distance increased but also identified several regions of the 12B1 assembly that were strain- or subclade-specific (Fig. S5). All strains maintained high sequence coverage across 12B1’s genic space, ranging from 97.5% in other A-type strains to 91.2% in strain UTEX 995 (Fig. 2B, Table S11). This is pattern is similar to that seen in interspecific comparisons of *Arabidopsis* spp. (Streptophyta), in which protein-coding genes are conserved while intergenic sequences show significant divergence (44).

### Phylogenetically divergent haplotypes within a single strain

Average breadth of coverage was strikingly low for most strains when aligned to the UTEX 2797 assembly (Fig. 2B, Table S11). The two exceptions were UTEX 2797 (*i.e*., aligned to itself, BOC = 86%) and another strain from Texas, 12A1 (BOC = 91.7%). This pattern of low coverage across the UTEX 2797 assembly is consistent with UTEX 2797 having two divergent haplotypes. Strain 12A1 likely shares these or closely related haplotypes as its reads map well to both haplotypes while reads from other strains primarily map to one haplotype or the other (Fig. 2B, Table S11). To investigate the evolutionary history of the UTEX 2797 haplotypes, we first plotted BOC across the 66 scaffolds of the UTEX 2797 reference genome. Clear haplotype blocks were identified that showed either high coverage in A-type strains from the U.S.A. or high coverage in the two A-type strains from Europe: CCMP2941 from Russia and RCC3703 from the United Kingdom (Fig. S6). Similarly, manual inspection of individual gene trees revealed that UTEX 2797 frequently grouped with these two European strains (*e.g*., Fig. S7) despite grouping with other strains from the U.S.A. in both the concatenation and coalescent species trees (Fig. 2A).

The BOC analysis suggested that the two haplotypes of UTEX 2797 have divergent evolutionary histories. This could lead to high levels of incongruence between the input gene trees (in which UTEX 2797 may group in two locations reflecting the evolutionary histories of its two haplotypes) and the inferred species tree (which forces each strain to appear only once). Therefore, we performed a gene tree-species tree reconciliation analysis using GRAMPA (45) to test whether a multi-labeled (MUL) species tree in which UTEX 2797 appeared twice was a more parsimonious representation of the input gene trees compared to the original single-labeled species tree. Seven of 29 possible MUL phylogenies had a smaller parsimony score, *i.e*., required fewer total gene duplication and loss events, compared to the single-labeled species tree (Fig. S8, Table S12). In the most parsimonious MUL species tree, UTEX 2797 grouped in two locations: with other A-type strains from the U.S.A (hereafter clade A1) and with CCMP2941 and RCC3703 from Europe (hereafter clade A2) (Fig. 3). Plotting the genomic distribution of UTEX 2797 genes that group with either clade A1 or A2 revealed a striking pattern. Most scaffolds formed syntenic pairs in the which one scaffold had a high proportion of genes that grouped with subclade A1 and the second scaffold contained genes that grouped with subclade A2 (Fig. 3, Table S13). To quantify the divergence between the UTEX 2797 haplotypes, we compared the distribution of synonymous substitutions of UTEX 2797 genes that grouped with either the A1 or A2 clades. This analysis revealed a bimodal distribution of *Ks* values in which A1 genes in UTEX 2797 were essentially identical to A1 strains 12B1 and CCMP3037 (*Ks* = 0) and more divergent to A2 strains RCC3703 and CCMP2941 (*Ks* = 0.009) (Fig. S9). This pattern was reversed in A2 genes (Fig. S9). From these results we can infer that UTEX 2797 was formed via the hybridization of an A1-like and an A2-like parent.

**Figure 3.**
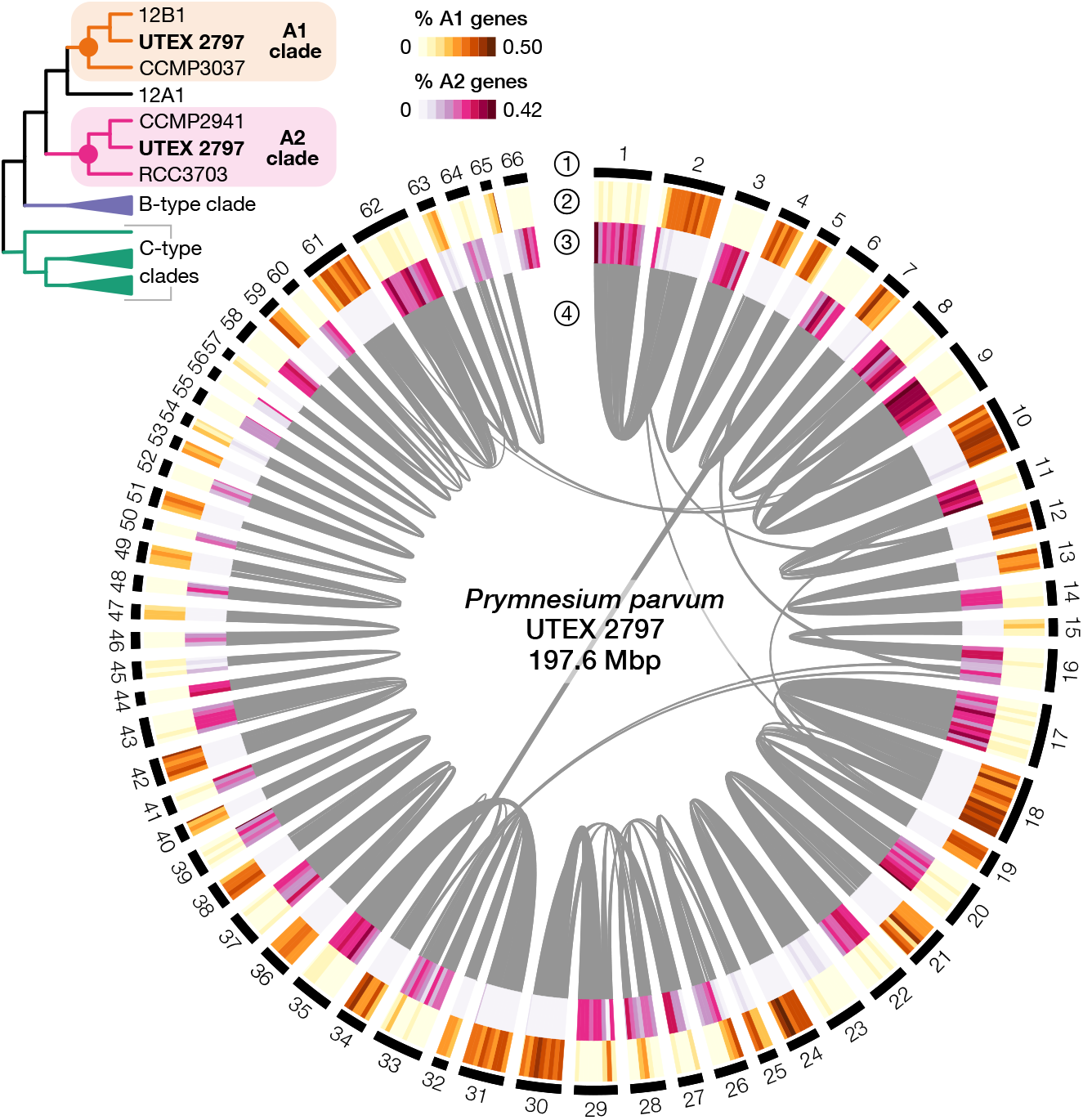
Phylogenetically distinct haplotypes of *P. parvum* strain UTEX 2797. Circos plot showing the 66 scaffolds of UTEX 2797 with four tracks (1) outer black track indicates scaffolds, (2) orange heatmap illustrates the percentage of genes in 50 kbp windows that group within the A1 clade, (3) pink heatmap indicates the percentage of genes in the same 50 kbp windows that group within the A2 clade, and (4) syntenic blocks (≥ 15 syntenic genes per block) are designated as gray bands. No branch support threshold was imposed for A1/A2 clade calls; the general pattern was the same but less vibrant when an ultrafast branch support threshold of 90 was imposed (Fig. S15). Multi-labeled (MUL) species tree (top left) shows the topology of the A1 and A2 clades as determined by the GRAMPA gene tree-species tree reconciliation analysis.

### Inter-strain variation in DNA content and haploid genome size

Flow cytometry revealed that DNA content varied dramatically among *P. parvum* strains (Fig. 4A). Strain 12B1 had the smallest amount of nuclear DNA at 115.6 Mbp, while K-0081 (a B-type strain) had the largest at 845.6 Mbp. Large variation in DNA content between strains of *P. parvum* has previously been interpreted as differences in ploidy (34). However, in our analysis, the relative DNA content between strains did not discretely cluster around integer fold changes (Fig. 4A), as would be expected if all variation in nuclear DNA content was due to differences in genome copy number (*i.e*., ploidy). Given the sequence-level divergence between strains in our analysis, it is possible that differences between strains could extend to the size of their haploid genome as well. Moreover, due to the limited understanding of the *P. parvum* lifecycle, the ploidy of different strains was not immediately clear. This creates a circular puzzle, as we cannot infer the haploid genome size of a strain from flow cytometry without *a priori* knowing the strain’s ploidy.

**Figure 4.**
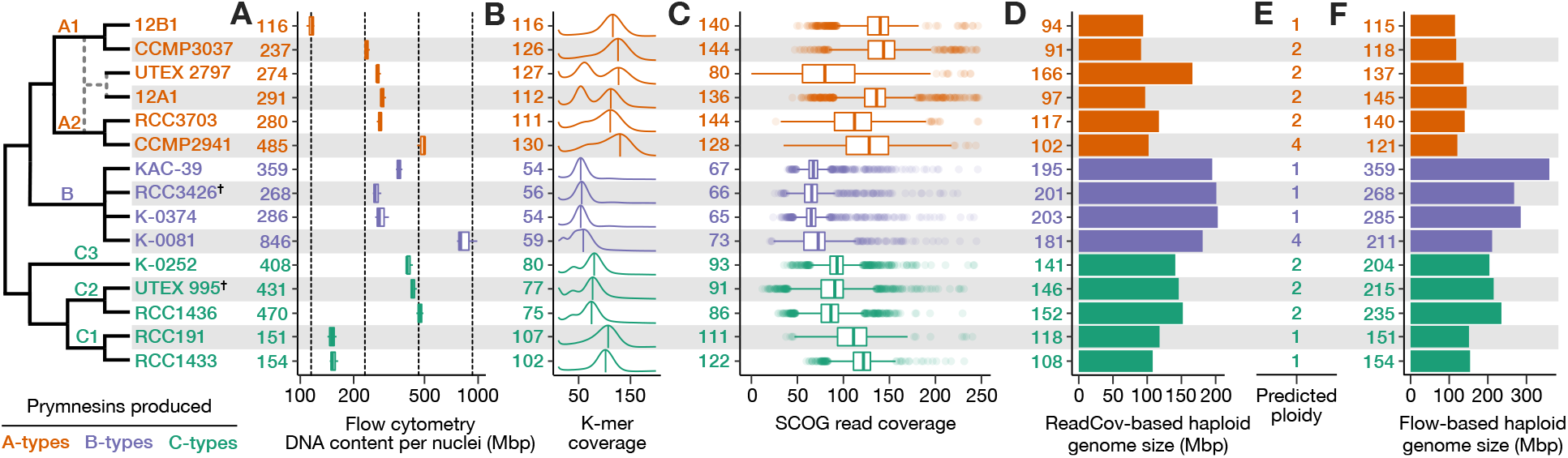
Summary of ploidy, heterozygosity, and genome size diversity in *P. parvum*. Evolutionary model (left) depicts strain relationships, including a predicted hybridization (dashed lines) giving rise to strains UTEX 2797 and 12A1. The chemotypes of strains RCC3426 and UTEX 995 (†) are inferred based on phylogenetic placement. A) Boxplots depicting total DNA content per nuclei (Mbp) based on flow cytometry. Numbers (left) indicate mean Mbp per strain. Vertical dashed lines indicate 1x, 2x, 4x, 8x Mbp relative to 12B1. B) K-mer coverage plots depict estimated heterozygosity; numbers indicate CMUKs, *i.e*., the homozygous k-mer peaks labeled by the labeled by vertical bars. Heterozygosity can be qualitatively assessed by the presence and relative height of a second peak at half the k-mer coverage of the homozygous peak. C) Boxplots depicting the distribution of read coverage for 2699 SCOGs; numbers indicate median read coverage. D) Haploid genome size was estimated using read coverage (total read length divided by median Illumina read coverage of SCOGs). E) Predicted ploidy was determined by cross-referencing DNA content (A) with sequencing-based estimated of genome size (B-D). F) Haploid genome size was estimated using flow cytometry (total DNA content divided by proposed ploidy).

To address this limitation, we used two sequence-based approaches to estimate the relative haploid genome size between strains. First, k-mer frequency analyses were performed using the Illumina reads from each strain. Reads were filtered to exclude possible contamination and depth normalized to permit inter-strain comparison of k-mer coverage. The coverage of maximal unique k-mers (CMUK) of the homozygous peak in k-mer frequency plots was used as a proxy for haploid genome size, with larger CMUKs indicative of smaller haploid genomes and vice versa. CMUK varied dramatically between strains with different prymnesin chemotypes, ranging from an average of 55.8 in B-type producers to 120.3 in A-types (Fig. 4B, Table S14). Average CMUK also varied between C-type strains with strains in the C1 clade having an average CMUK of 104.5 compared to strains in C2 and C3 clades, which had and average CMUK of 77.3 (Fig. 4B, Table S14). This pattern indicates considerable differences in haploid genome size between clades with the A-type clade having the smallest haploid genome size and the B-type clade having the largest.

The amount of heterozygosity also varied considerably between strains (Fig. 4B). In addition to 12B1, six strains (A1 strain CCMP3037; B strains K-0374, KAC-39, RCC3426; and C1 strains RCC191, RCC1433) showed little to no evidence of heterozygosity, which could indicate that, like 12B1, these strains are haploid. However, in the case of A1 strain CCMP3037, its flow cytometry estimated DNA content was twice that of 12B1 despite the two strains having similar CMUKs peaks (Fig. 4A,B). This indicates that CCMP3037 is a diploid strain with low levels of heterozygosity. For all other low heterozygosity strains in B and C1 clades, decreased CMUK relative to 12B1 corresponds with increased DNA content, suggesting that these strains are haploids. All other strains displayed comparatively moderate to high levels of heterozygosity, indicating that they are either diploids (2n) or polyploid (3n or greater). Notably, strain 12A1 had a pronounced heterozygous peak indicating extremely high levels of heterozygosity, similar to that observed in UTEX 2797 (Fig. 4B). This, coupled with 12A1’s high BOC against the UTEX 2797 diploid assembly, indicates that 12A1 is also a hybrid strain.

Average read coverage across 2699 single-copy orthogroups (SCOGs) was used as a second sequence-based estimate of haploid genome size (Fig. 4C, Table S14). SCOG and CMUK coverage estimates were in agreement for most strains, except for UTEX 2797, which had significantly lower average SCOG coverage compared to the location of its CMUK peak (Fig. 4B,C). This discrepancy is due to the nature of its hybrid genome and Hi-C scaffolded assembly with resolved haplotypes. To be assigned to a SCOG, genes in UTEX 2797 have likely returned to single-copy post hybridization, resulting in lower coverage compared to genes that have been retained in duplicate. We used the Lander-Waterman equation to estimate the haploid genome size for all strains based on the total length of the Illumina sequencing library divided by the median SCOG coverage, which served as an estimate of coverage for the entire genome (46, 47). This sequence-based estimate of haploid genome size ranged from 91 Mbp in A-type strain CCMP3031 to 203 Mbp in B-type strain K-0374 (Fig. 4D).

Cross-referencing total DNA content with the sequence-based haploid genome size, we were able to assign a predicted ploidy level for each strain (Fig. 4E, Table S14). Six strains were determined to be haploids, and seven (including hybrids 12A1 and UTEX 2797) were diploids. Two strains, A-type CCMP2941 and B-type K-0081 appear tetraploid. Using this predicted ploidy, we also determined flow cytometry-based estimates of haploid genome size (Fig. 4F, Table S14). The A clade has the smallest genome size (average = 130 Mbp), followed by the C1 clade (153 Mbp). The C3 strain (K-0252) had an estimated genome size of 204 Mbp, and the C2 clade had an average genome size of 225 Mbp. Lastly, B clade strains had the largest genomes (average = 281 Mbp), while also having the largest amount of intra-clade variation in genome size (Fig. 4F).

We investigated the relationship between DNA content and average cell volume and did not identify a clear trend (Fig. S10). When comparing between haploid strains, we found a weak association between cell size and prymnesin type, with B-types having the largest genomes and largest cells and the single A-type haploid (12B1) having both the smallest genome and cell size (Fig. S10B). However, we did not find a similar correlation when comparing between diploid strains. When looking within prymnesin A-type producing strains, we found a weak positive correlation between cell size and ploidy state, but we did not find a similar association within B- or C-type strains (Fig. S10C).

### Accessory gene families and horizontal gene transfer

To investigate the impact of genome variation on gene family evolution in *P. parvum*, we performed a pangenome analysis of OrthoFinder-predicted gene families and identified a total of 47,043 orthogroups including 16,453 core orthogroups present in all strains; 20,738 accessory orthogroups present in 2-14 strains; and 9852 singleton orthogroups unique to a single strain (Table S15). The number of shared orthogroups generally clustered by prymnesin type, with some exceptions (Fig. 5A; Table S16). Strains 12B1 and UTEX 2797 did not cluster with other A-types, likely in part due to the differences between their Hi-C scaffolded assemblies and the other Illumina-only assemblies. The C-types also clustered into two groups based on orthogroup membership, with C2 strains (RCC1436 and UTEX 995) clustering separate from C1 and C3 strains (Fig. 5A).

**Figure 5.**
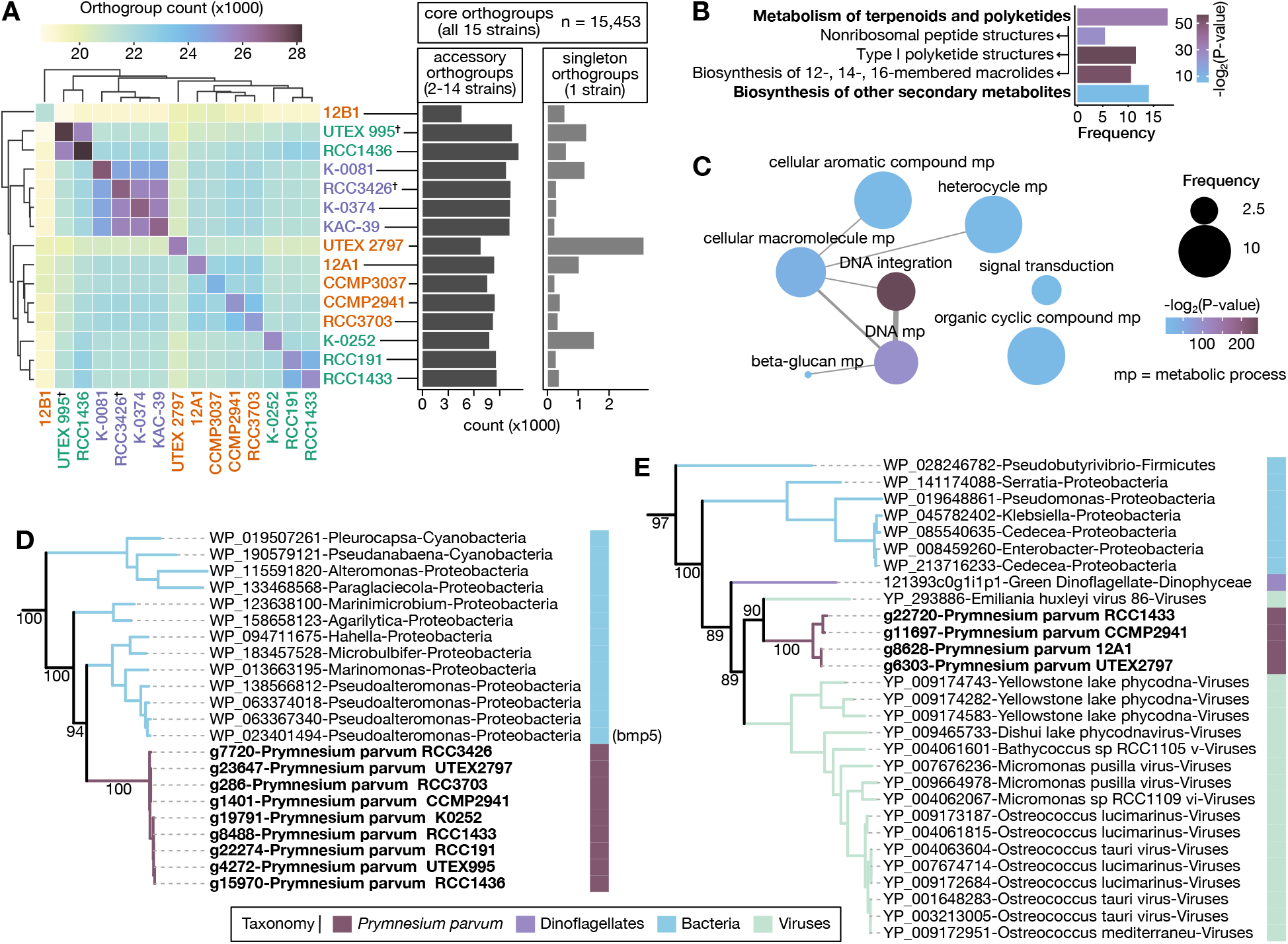
Pan genome analysis and horizontal gene transfer in *P. parvum*. A) Hierarchically clustered heatmap showing the number of orthogroups shared by each strain pair. Strains are colored based on the prymnesin type produced (as in Fig. 2). The chemotypes of strains RCC3426 and UTEX 995 (†) are inferred based on phylogenetic placement. Center diagonal indicates the total number of orthogroups, including singletons, present in each strain. Bar charts indicate the number of accessory orthogroups and singleton orthogroups in each strain. B) Significantly enriched KEGG pathways (unbolded) and pathway categories (bolded); arrows point to KEGG pathway parent category. Bar height indicates frequency of the annotation in accessory orthogroups with one or more KEGG annotations. C) Significantly enriched GO categories in accessory orthogroups. Width of network edges indicate the degree of similarity between GO terms as calculated by REVIGO (63). Bubble size indicates frequency of the annotation in accessory orthogroups with one or more GO annotations. See Table S17 for list of all functional enrichment tests performed. D) Flavin-dependent halogenase phylogeny showing HGT from marine bacteria. The *bmp*5 gene functionally characterized in *Pseudoalteromonas* is labeled. E) Clp protease phylogeny showing HGT from viruses. For both phylogenies, numbers along select branches indicate ultrafast bootstrap support values for the descendant nodes. Support for all branches and expanded phylogenies are available in Fig. S11.

A significant percentage of all orthogroups (44.1%) were variably present in 2-14 strains. These accessory orthogroups belong to several enriched functional categories that could be associated with metabolic and genome size variation in *P. parvum*. Several KEGG specialized metabolic pathways, including those for the biosynthesis of type I polyketides, macrolides, and nonribosomal peptides, were enriched in accessory orthogroups compared to core and singleton orthogroups (BH adjusted p-value < 0.01; Fig. 5B, Table S17). Moreover, 31 Gene Ontology (GO) categories were enriched in the accessory orthogroups, the most significantly enriched of which was GO:0015074, DNA integration (BH adjusted p-value = 3.68E-73; Fig. 5C, Table S17). Most orthogroups assigned to this GO category were annotated as integrase-like enzymes, common components of lysogenic viruses and transposable elements. We investigated the 60 genes annotated with the DNA integration GO term (GO:0015074) in 12B1 and UTEX 2797 and found that all genes fell within 400 bp of a predicted repeat. Many of these genes were found within Ngaro LTRs, which intersected 31.3% of integrase-like genes in 12B1. This pattern reveals that enrichment of the DNA integration GO term is driven by transposable element integrase genes that have been incorporated into the gene model predictions. We investigated the phylogenetic distribution of this pattern by performing functional enrichment on gene families uniquely present in A-, B-, and C-type strains. We found that the DNA integration GO term was enriched in the B- and C-type specific orthogroups (BH adjusted p-values = 3.94e^−15^ and 1.72e^−4^, respectively) but was not significantly enriched in A-type specific orthogroups (Table S17), suggesting that an expansion of transposon copy number may contribute to the greater haploid genome sizes of the B- and C-type clades.

One potential source of accessory genes in a pangenome is horizontal gene transfer (HGT) (48). To investigate the role of HGT in the genome evolution of *P. parvum*, we performed an Alien Index (AI) analysis (49) and focused our investigation on HGTs into *Prymnesium* after the genus diverged from other haptophytes. The AI screen flagged 95 orthogroups as candidate HGTs (Table S18). We used a custom phylogenetic pipeline to manually evaluate all candidate HGTs and found that most (n = 61) were phylogenetically inconclusive or lacked support for HGT. Of the 34 HGT candidates that passed manual inspection, we focused on eleven HGTs with clear donor lineages with strong node support (Fig. S11, Table S19). Two HGTs were likely acquired from eukaryotic donors: HGT01, a bicarbonate transporter of diatom origin, and HGT02, a gene of unknown function that grouped with pelagophytes. Eight HGTs (HGT03 - HGT10) were likely acquired from bacteria. All the bacterially derived HGTs were enzyme-coding with diverse metabolic activities (Table S19). For example, HGT09 grouped phylogenetically with sequences from marine bacteria (Fig. 5D), including *bmp5*, a decarboxylating flavin-dependent halogenase involved in the biosynthesis of polybrominated natural products in *Pseudoalteromonas* spp. (50). Lastly, HGT11 was a Clp protease that grouped with large dsDNA megaviruses that infect eukaryotic algae (Fig. 5E).

Several horizontally transferred genes were part of the accessory genome and variably present across strains (Table S19). Some of this variation is likely due to ancestral HGTs being lost in some clades; *e.g*., HGT07 and HGT10 both appear to have been gained in an ancestor of *Prymnesium* and subsequently lost in the B-type clade (Fig. S11). HGT11, the Clp protease gene of viral origin, was notable due to its presence in only four disparate strains: three A-type strains (UTEX 2797, 12A1, and CCMP2941) and one C-type (RCC1433) (Fig. 5E). If this gene was acquired in a shared ancestor of all four strains, at least six independent loss events would be required to explain this presence/absence pattern. An alternative explanation could be that multiple independent acquisitions of a viral Clp protease have occurred in *P. parvum*. The genomic neighborhood of HGT11 in UTEX 2797 contains six additional genes with a top BLAST hit to EhV-86, a dsDNA megavirus that infects the haptophyte *Emiliania huxleyi* (51). These additional genes were not recovered in our primary screen for HGT due to a limited number of hits to proteins in the NCBI RefSeq database, which were insufficient for phylogenetic analysis (see SI Materials and Methods). However, their sequence similarity suggests that a block of viral genes was likely acquired in a single HGT event in the lineage leading to UTEX 2797 (Fig. S12). We checked for shared synteny between UTEX 2797 and the three additional strains that contain HGT11. In 12A1, HGT11 is located on a 20-gene scaffold (Scaf652979) which shares nearly perfect synteny with UTEX 2797, including five genes within the proposed viral fragment (Fig. S13), which suggests that UTEX 2797 and 12A1 share the same HGT event. In contrast, the scaffolds that contain HGT11 in CCMP2941 and RCC1433 are syntenic with each other, but neither are syntenic with UTEX 2797 (Fig. S13). This suggests that the HGT that gave rise to the Clp protease in CCMP2941 and RCC1433 was likely independent from that of UTEX 2797 and 12A1. Long-read assemblies of additional strains including 12A1, CCMP2941, and RCC1433 are needed to confirm this pattern.

## DISCUSSION

The harmful algal bloom-forming eukaryote, *Prymnesium parvum*, possesses considerable genomic diversity. Combining data from synonymous substitutions, reference genome coverage, phylogenetics, and genome size, the fifteen strains in our analysis can be subdivided into six distinct clades consisting of at least three cryptic species (Fig. 4). Evidence for these cryptic species comes from the fact that prymnesin chemotype is phylogenetically segregated, which indicates that the *P. parvum* A-, B-, and C-type clades are reproductively isolated. Here, we provide further support for these clades being separate cryptic species based on the extreme variation in their genome size, as excessive chromosome-level differences will likely inhibit proper chromosome pairing during meiosis. The most dramatic difference in genome size occurred between the sister clades of A- and B-type strains (Fig. 4F). These clades show limited substitutions at synonymous sites (*Ks* = 0.02; Fig. 2A), which suggests that the changes to their genome sizes have been relatively recent. Possible mechanisms for genome expansion in B-type strains includes whole-genome duplication (WGD) and/or proliferation of transposable elements. Our analysis of *Ks* distributions showed no evidence of recent WGD in the last common ancestor of the B-clade (Fig. S4). Instead, gene functions associated with transposable elements were enriched in gene families unique to B-type strains (Table S17), suggesting that the increase in their genome size could be due to an increase in transposition activity of these elements. A high-quality reference genome of a B-type strain would enable further investigation of repeat diversity and expansion across these two clades and clarify the mechanism(s) of genome size variation in *P. parvum*.

Although *P. parvum* is currently considered a single morphospecies, earlier taxonomic descriptions split the species in two (*P. parvum* and *P. patelliferum*) based on slight differences in the morphology of the organic scales that cover their cell surface (52). When the rDNA ITS1 region was found to be identical in strains of *P. parvum* and *P. patelliferum* isolated from the same geographic location, the two species were merged into one (53). Later, two strains originally identified as *P. parvum* (K-0081 and strain RL10parv93, not included in our analysis) were found to have higher amounts of DNA compared to three strains labeled *P. patelliferum* (K-0252, RCC191, and a third strain not assessed here, RLpat93) (34). This observation led Larsen and Edvardsen (1998) to propose a cryptic lifecycle for *P. parvum* that alternates between flagellated haploid and diploid forms. Our analysis identified strains of different ploidy states, supporting the existence of a cryptic sexual lifecycle. Moreover, the existence of a hybrid strain (UTEX 2797) with two phylogenetically divergent parental genotypes indicates that syngamy can occur in *P. parvum*. Our assessment of ploidy across these taxa, which includes flow cytometry and read-based estimates of haploid genome size, indicates that K-0081 is a tetraploid, not a diploid as previously thought, and that K-0252 is a diploid and not a haploid (Fig. 4). Given the inclusion of these strains in the hypothesis about scale morphology and ploidy, our new results suggest that scale morphology is not diagnostic of ploidy state. Analysis of the fifteen strains reported here places variation in ploidy and genome size in a phylogenetic framework, facilitating future investigation into cellular morphology and sexual reproduction in *P. parvum*.

Our phylogenomic analysis also reveals that UTEX 2797, a Texas *P. parvum* strain frequently used in growth and toxicity experiments, is a hybrid of A1 and A2 parents (Fig. 4A). It is intriguing that the A1 strains in our analysis are from the U.S.A while the A2 strains are from Europe, and it is tempting to speculate that hybridization was a result of a recent introduction of the A2 lineage in Texas. Our analyses suggest that the high heterozygosity strain 12A1 is a hybrid as well, but it is unclear from our analysis if 12A1 arose from the same hybridization event as UTEX 2797. Further investigation into the origin of these strains requires additional taxon sampling of both A1 and A2 lineages as well as long-read based assemblies of 12A1 to resolve its two haplotypes. The evolutionary outcome of hybridization in *P. parvum* is also unknown. Are these hybrid strains capable of sexual reproduction? Genome divergence between A1 and A2 clades may be such that homologous chromosomes are unable to pair correctly during meiosis. If so, these hybrids may be effectively trapped as diploids and only able to reproduce asexually. Alternatively, whole-genome duplication following hybridization (allopolyploidy) has been shown to restore fertility in yeast (54), which could make these *P. parvum* hybrids reproductively haploids. Characterizing the sexual lifecycle of *P. parvum* would allow researchers to address several outstanding questions regarding the ecological and evolutionary consequences of extreme genetic variation in these toxic bloom forming eukaryotes. Knowledge of the *P. parvum* lifecycle would enable the design of mating tests to determine the reproductive status of hybrid strains. Mating tests could also be used to assess reproductive isolation between divergent populations, as has been done in other morphologically indistinguishable cryptic species complexes (55–57).

Previous work indicates that HABs of *P. parvum* are comprised of multiple genotypes (21, 24, 58), and our results reveal that the genome-level differences between these genotypes can be dramatic. For example, strains 12A1 and 12B1 were isolated from the same Texas bloom in 2010 and consistently show different cell-level behaviors involved in toxicity and predation of microalgal prey (21, 24). Here, we show that characteristic differences between 12A1 and 12B1 extend to large differences in their genomes as well, with 12A1 being a hybrid diploid and 12B1 having a streamlined haploid genome. Moreover, significant gene-level differences exist between these two strains, with 7% (n = 1485) of 12B1 orthogroups absent in 12A1. An even larger percentage, 22% (n = 5709), of 12A1 orthogroups are absent in 12B1 (Table S16), including one that was horizontally acquired (HGT11; Fig. 5E). Gene families that are variably present across strains in our analysis include those for the biosynthesis of type I polyketides, (Fig. 5B), the class of specialized metabolites that includes prymnesins (17). This is unsurprising given the structural diversity in prymnesins that have already been characterized in *P. parvum* (18, 19). However, it is notable that 12A1 and 12B1, both A-type strains, show variable representation of eighteen orthogroups assigned to the type I polyketide KEGG pathway (map01052; Table S17, Table S15). If any of these orthogroups are involved in toxin biosynthesis, phenotypic differences between these two strains could extend to their toxin profiles as well. The coding capacity of *P. parvum* has also been expanded by HGT. Prymnesins are halogenated metabolites, and though we have not functionally characterized it, it is tantalizing to note that a halogenase is among the HGT candidates in the *P. parvum* genomes (HGT09; Fig. 5D). HGT is often a source of metabolic innovation (*e.g*., 49, 59–61) and elucidating the prymnesin pathway will determine if this gene is relevant to the production of these metabolites. More work is needed to identify the genetic factors associated with toxin production and the phenotypic differences between these diverse strains of *P. parvum*. Additionally, work is needed to identify the selective advantage of different phenotypes and whether such advantages fluctuate between bloom and non-bloom conditions. The genome assemblies and phylogenomic analysis reported here are thus essential resources that will enable future investigation into the eco-physiological consequences of hidden genetic diversity in this HAB-forming morphospecies.

## MATERIALS AND METHODS

Strains and their respective media types and culturing conditions are summarized in Table S8. Flow cytometry for genome size estimation and LC-MS for prymnesin chemotyping were performed as described in SI Materials and Methods. Details on genome sequencing and assembly, repeat and gene prediction, and assessment of assembly completeness are described in SI Materials and Methods. Analysis of synteny within the UTEX 2797 Hi-C scaffolded genome assembly and between assemblies of UTEX 2797 and other *P. parvum* strains was performed using the online Comparative Genomics Platform (CoGe) as described in SI Materials and Methods. Orthologous gene families, individual gene trees, concatenation and coalescent-based species trees, and gene tree-species tree reconciliation was performed using the phylogenomics workflow summarized in Fig. S14 and described in SI Materials and Methods. Breadth of coverage (BOC) was calculated from alignments of subsampled, filtered Illumina reads to the 12B1 and UTEX 2797 scaffolded assemblies. See SI Materials and Methods for details on read filtration to remove contamination and subsampling to control for library differences between strains. Depth of coverage was calculated on a per-base level and BOC was calculated as the proportion of base-pairs in the final assembly that had coverage greater than N coverage. BOC using coverage N = 0, 5, 10, 20, and 50 were all evaluated. Final BOC tables for each assembly can be found in Tables S11, and additional details are provided in in SI Materials and Methods. Heterozygosity, coverage of maximal unique k-mers, and SCOG read coverage were estimated using the subsampled, filtered Illumina gDNA reads as described in SI Materials and Methods. The haploid genome size (G) was calculated using the Lander-Waterman equation (46): *G* = *LN*/*C* where LN is the total combined length (in base pairs) of the input Illumina reads and C is coverage as estimated by SCOG read depth. All additional method details including cell size estimation, functional annotation, tests for functional enrichment, and detection of horizontal gene transfer are described in SI Materials and Methods.

## DATA AVAILABILITY

All genome assemblies, predicted CDS and protein sequences, multiple sequence alignments, tree files, and other related data files are available through FigShare (https://doi.org/10.6084/m9.figshare.21376500). Scripts are available through GitHub (https://github.com/WisecaverLab/Pparvum_genome_diversity). Raw sequencing reads have been deposited in the Sequence Read Archive database under accession number PRJNA807128. LC-MS data from chemotyping is available in the EBI MetaboLights database under accession MTBLS5893. Strains 12B1 and 12A1 have been deposited in the UTEX Culture Collection of Algae at UT-Austin under UTEX accessions UTEX LB ZZ1299 and UTEX LB ZZ1300).

## ACKNOWLEDGEMENTS

We thank members of the Wisecaver lab, Dr. Jody Banks, and Dr. Brian Dilkes for helpful discussions as well as Dr. Clint Chapple and Dr. Scott McAdam for plant tissue for flow cytometry. This work was conducted in part using the resources of the Rosen Center for Advanced Computing at Purdue University and well as through use of the Flow Cytometry and Cell Separation Facility at the Bindley Bioscience Center at Purdue, a core facility of the NIH-funded Indiana Clinical and Translational Sciences Institute. This work was supported by the National Institute for Environmental Health Sciences under grants F32-ES032276 to TRF and R21-ES032056 to BSM and the National Science Foundation under grant DEB-1831493 to JHW and WWD.

