## Supplemental text and figures for "Extreme genome diversity and cryptic speciation in a harmful algal bloom forming eukaryote"

### Supporting Information Text

#### SI MATERIALS AND METHODS

**Culturing methods**. Strains and their respective media types are summarized in Table S8. Cultures were kept at 20 °C using a 12:12 light dark cycle and irradiance of 40-200 µmol photos $m^{-2}$ $s^{-1}$.

**Flow cytometry**. Nuclei for genome size estimation were isolated using LB01 buffer (15 mM Tris, 2 mM $Na_2$-EDTA, 0.5 mM spermine tetrahydrochloride, 80 mM KCl, 20 mM NaCl, 0.1% (v/v) Triton X-100, pH=8.0) (1, 2). For *P. parvum,* 1 mL of exponentially growing culture was centrifuged at 1000 x g for 10 minutes. The supernatant was decanted, and the cell pellet was flash-frozen in liquid nitrogen. The pellet was resuspended in 1 mL of ice-cold LB01 buffer. For the *Chlorella vulgaris* genome standard, 1 mL of exponentially growing liquid culture (Carolina Biological) was centrifuged at 1000 x g for 10 minutes. The cell pellet was resuspended in 1 mL ice-cold LB01 buffer and transferred to a 2 mL ZR BashingBead Lysis Tube (Zymo Research Cat #S6003-50). The tube was agitated using a Disruptor Genie (Scientific Industries Model #SI-D238) for 5 minutes. For the *Selaginella moellendorfii* genome standard, leaf and stem tissue (provided by Dr. Scott McAdam, Purdue University) were chopped using a razor blade with 1.5 mL ice-cold LB01 buffer added per 100 mg tissue for 3 minutes. For the *Arabidopsis thaliana* genome standard, rosette tissue from the Col-0 reference accession (provided by Dr. Clint Chapple, Purdue University) was chopped using a razor blade with 1.5 mL ice-cold LB01 buffer added per 100 mg tissue for 3 minutes. All nuclei suspensions were filtered through a 40 µm cell strainer and kept at 4 °C until use.

RNaseA and propidium iodide were added to nuclei suspensions at a final concentration of 0.4 mg/mL and 0.05 mg/mL respectively. After briefly vortexing, the nuclei were incubated at 4 °C in the dark for 3 hours. Samples were analyzed using a BD Accuri C6 Plus flow cytometer. All samples were gathered using a flow rate of 14 µL/min and a core size of 10 µm. Fluorescence values were gathered using a 488-nm laser using a 585/40 nm band pass filter and a 670 nm long pass filter. Data from at least 300 nuclei were collected per sample. Biological replicates were performed on three subsequent afternoons at approximately the same time. Samples were processed using the FlowCal Python library (3). A line of best fit through the genome standards was calculated using ordinary least squares linear regression (Fig. S16). The linear model was then used to estimate the nuclear DNA content in picograms for *P. parvum* strains.

**Prymnesin chemotyping.** *P. parvum* strains were cultured in media consisting of L1 (Guillard and Morton, 2003) in GF/F glass fiber filter (Whatman, USA) filtered seawater (Scripps Institution of Oceanography seawater system) diluted to 25% salinity with 18 MOhm water with 1.5 mM added $NaHCO_3$ to adjust for the lack of carbon in 18 MOhm water (4). Cultures were grown in 100 mL of media in 500 mL glass Erlenmeyer flasks without external aeration or shaking. Exponential phase culture (>= $100e^3$ cells/mL, <= $600e^3$ cells/mL; late log phase) was harvested via filtration onto 47 mm glass fiber filters (GF/B of GF/F, Whatman) using laboratory vacuum. The resulting filters were stored in 50 mL polypropylene (PP) centrifuge tubes at -80 °C until extraction.

PP tubes with the collected biomass filters were shaken with 25 mL of ethyl acetate at 37 °C (220 RPM) for 20 minutes, and the resulting yellow-green extract was discarded. We chose ethyl acetate rather than the standard cold acetone for pre-extraction based on reports that the cold-acetone lipid-solubilization step can lead to PRYM losses due to variable quantities of water on the filter and resulting extraction (4). The ethyl acetate pre-extraction step was repeated 2-3 times until the resulting extract was without coloration. The tube was shaken with 25 mL of methanol (MeOH) at 37 °C for 30 minutes. After centrifugation at 4000 x g for 10 minutes at 20 °C, the MeOH extract was decanted into a 25 mL pear-shaped flask (P/N 9477-06, Ace Glass) and rotary evaporated to dryness at 30 °C under reduced pressure (50 mbar, 150 RPM). The resulting residue was redissolved in 500 µL MeOH and filtered through a 0.2 µm PFTE filter (P/N: CIPT-02, American Chromatography Supplies) into a glass HPLC vial. For each sample, 20 µL was injected onto an Agilent 1260 Infinity HPLC system coupled to an Agilent 6530 quadrupole time-of-flight (QToF) mass spectrometer.

Compounds were separated via gradient HPLC on an Agilent 1260 Infinity system, via C18 reversed phase chromatography (Phenomenex Kinetex C18 150 x 4.6 mm, 5 μm, 100 Å, with C18 guard column), at 0.7 mL/min with the following solvents and gradients: Solvent A: $H_2O$ + 0.1% v/v formic acid, Solvent B: acetonitrile + 0.1% v/v formic acid. Compounds were eluted with a gradient of 10% to 100% B over 30 minutes. To wash and equilibrate the column for subsequent injections, the gradient was then held at 100% B until 36 minutes, decreased to 50% over 3 minutes, returned to 10% over 3 minutes, and equilibrated at 10% B for 3 minutes for a total run time of 45 minutes. Under these conditions PRYM-1 and PRYM-2 eluted at around 55-57% B (15.19 to 15.68 min).

The Agilent 6530 QToF MS was configured in either the 3200 *m/z*, HiRes (4GHz) instrument state or the high dynamic range (2GHz) instrument state. In either case the instrument was quick tuned and mass calibrated just before use. The instrument was set to positive ionization mode. The source parameters were 300 °C gas temp, 11 L/min drying gas, nebulizer at 45 psig, source voltage at 4000V, fragmentor at 100V, Skimmer at 65V, OCT 1 Rf Vpp at 750 V. $N_2$ gas was supplied by a Parker NitroFlowLab $N_2$ generator at ~>90% purity. The reference mass infusion and locking were disabled, due to an overlap with the PRYM aglycone $[M+2H]^{2+}$ isotopic peaks with the vendor supplied lock mass. The LC flow was diverted to waste from 0 to 6 minutes and MS acquisition was not performed. At 6 minutes, the LC flow was switched to the MS source & the instrument began acquiring $MS^1$ data from 125-3200 *m/z*, at a rate of 4 spectra/second, in the auto $MS^2$ mode with a 250 ms dwell time per subsequent $MS^2$ scan. Between 1-3 of the most abundant precursors per $MS^1$ cycle were chosen for fragmentation, with dynamic exclusion of precursors after their selection. A fixed collision energy of 20 was used. All data was acquired in profile mode. MS acquisition was stopped after 43 minutes.

The resulting LC-MS data was transformed from Agilent MassHunter .d format to .mzML format using Proteowizard version 3.0.20303 (5), with parameters '--zlib'. Data in .mzML format were analyzed using with MZmine2 v2.53 (6), while .d format data were analyzed with Agilent MassHunter (B.05.00). PRYM-1 and PRYM-2 were identified by comparison of their characteristic multi-chlorinated $MS^1$ isotopologue ionization intensity pattern to *in silico* calculated $MS^1$ isotopologue intensity patterns using the enviPat Web 2.4 tool (7). Coeluting $[M+H]^+$, $[M+2H]^{2+}$, and presumed aglycone in-source fragments, each showing the characteristic $MS^1$ pattern, were detected.

**Genome sequencing and assembly**. Genomic DNA for Illumina sequencing was extracted from *P. parvum* cell pellets using the CTAB method according to the following protocol https://dx.doi.org/10.17504/protocols.io.b5qhq5t6 (8). Extracted DNA was purified using a Genomic DNA Clean and Concentrator kit (Zymo Research). Sequencing libraries were constructed and sequenced to produce 150 bp paired-end reads using one of two approaches: 1) libraries were prepared using a TruSeq DNA PCR-Free library prep kit (Illumina, San Diego, CA), and sequenced using an Illumina NovaSeq 6000 at the Purdue Genomics Center 2) libraries were prepared using an NEBNext DNA library prep kit (New England Biolabs Inc.) and sequenced using an Illumina NovaSeq 6000 by Novogene Corporation Inc. (Sacramento, CA). Illumina gDNA read quality was assessed by FastQC v0.10.0 (9). Short-read only genome assemblies were performed by Abyss v2.2.4 (10) using a k-mer size of 96. Contigs less than 500 bp in length and those flagged as bacterial contamination (see below) were discarded.

Bacterial contamination in the Illumina assemblies was identified using Blobtools v1.1.1 (11). For each strain, Illumina gDNA reads were aligned to the Abyss assembly using BWA-MEM v0.7.15 (12) to generate a coverage BAM file. Contigs were queried against the NCBI nucleotide (nt) database (accessed September 11, 2021) using blastn v2.11.0 (13). DIAMOND v2.0.8.146 (14) was used to query contigs against a custom protein databases that consisted of NCBI RefSeq (release 98) (15) sequences supplemented with additional predicted protein sequences from MMETSP (16) and the 1000 Plants transcriptome sequencing project (1KP) (17). The custom protein database used in the BlobTools analysis is available from the authors as well as through the following link: https://www.datadepot.rcac.purdue.edu/jwisecav/custom-refseq/2021-08-02/. The blobtools taxrule 'bestsumorder' determined the taxonomic assignment of each contig, prioritizing information from protein hits first. Contigs denoted as non-eukaryotic in origin were removed to produce the final filtered assembly. Lastly, BBSplit v38.87 (18) was used to retain only Illumina reads that mapped to the Blobtools filtered assembly (hereafter referred to as filtered Illumina reads). See Fig. S17 for example blobplots.

For long-read sequencing with Oxford Nanopore Technologies (ONT), high molecular weight DNA was extracted from isolated *P. parvum* nuclei using the following protocol https://dx.doi.org/10.17504/protocols.io.7b7hirn (19). At least 1.5 µg of gDNA was used as input for an Oxford Nanopore LSK-109 library ligation kit and sequenced on R9 MinION flow cells. Base calling was performed with Guppy v2.3.5 (Oxford Nanopore Technologies). Reads less than 3 kbp long or with quality scores less than 7 were discarded. Different assembly approaches were selected to optimize for either assembly contiguity (in the case of low heterozygosity 12B1) or the amount of resolved haplotypes (in the case of high heterozygosity UTEX 2797). The 12B1 long-read assembly was created using both Nanopore and Illumina gDNA data via MaSuRCA v3.3.1 (20) with the following parameters: LHE_COVERAGE=60, CA_PARAMETERS=cgwErrorRate=0.15, KMER_COUNT_THRESHOLD=2, CLOSE_GAPS=1, JF_SIZE=5000000000. The UTEX 2797 long-read assembly was created using ONT reads only via Canu v2.1.1 (21) with an expected genome size of 200 Mbp. Both assembly types were error corrected via five rounds of polishing with Illumina gDNA reads that were first aligned to the assembly with using BWA-MEM v0.7.15 (12) and polished with Pilon v1.23 using default settings (22).

Chromatin conformation capture data was generated using a Phase Genomics (Seattle, WA) Proximo Hi-C 2.0 Kit, which is a commercially available version of the Hi-C protocol (23). Following the manufacturer's instructions for the kit, intact cells were crosslinked using a formaldehyde solution, digested using the DPNII restriction enzyme, end repaired with biotinylated nucleotides, and proximity ligated to create chimeric molecules composed of fragments from different regions of the genome that were physically proximal *in vivo*. Molecules were pulled down with streptavidin beads, processed into an Illumina-compatible sequencing library, and sequenced on the Illumina HiSeq platform as 2x150 bp reads. Illumina reads were aligned to the long-read assemblies (Canu assembly for UTEX 2797 and MaSuRCA assembly for 12B1) using BWA-MEM v0.7.15 (12) with the -5SP options specified, and all other options default. SAMBLASTER (24) was used to flag PCR duplicates, which were then excluded. Alignments were filtered with samtools (25) using the -F 2304 filtering flag to remove non-primary and secondary alignments. Putative misjoined contigs were broken using Juicebox (26, 27) based on Hi-C alignments. Kraken v2 (28) identified eukaryotic contigs, which were selected for scaffolding, and prokaryotic contaminants, which were discarded. The same alignment procedure was repeated from the beginning on this corrected assembly. Phase Genomics' Proximo Hi-C genome scaffolding platform was used to create chromosome-scale scaffolds from the corrected assembly as previously described (29). As in the LACHESIS method (30), this process computes a contact frequency matrix from the aligned Hi-C read pairs, normalized by the number of DPNII restriction sites (GATC) on each contig, and constructs scaffolds that optimize expected contact frequency and other statistical patterns in Hi-C data. Approximately 60,000 separate Proximo runs were performed to optimize the number of scaffolds and scaffold construction to make the scaffolds as concordant with the observed Hi-C data as possible.

During the Alien Index analysis (see Horizontal Gene Transfer methods section below), we flagged two 12B1 MaSuRCA contigs (scf7180000001543 on 12B1-Scaf8 and scf7180000001202 on 12B1-Scaf32) as bacterial contamination in the scaffolded 12B1 assembly. Both contigs were located at the ends of scaffolds, all genes on the contigs were of bacterial origin, and the percent identity to the database sequences was high (> 80% for all genes), indicating that these bacterially derived contigs were incorporated into the Hi-C scaffolded assembly in error rather than true horizontal gene transfer. Therefore, these contig sequences were manually removed from the final genome assembly and the flagged genes filtered from the final gene set. The resulting 12B1 assembly and gene annotations following these steps were designated as final (v1).

**Repeat and gene prediction.** *De novo* repeat identification was separately performed on the scaffolded assemblies of strains 12B1 and UTEX 2797 using RepeatModeler v2.0.1 (31). The resulting modeled libraries were used to inform repeat masking the assemblies with RepeatMasker v4.0.7. Repeats were also masked using the UTEX 2797 *de novo* repeat library for the short-read only assemblies.

To maximize capture of the *P. parvum* transcriptome for gene calling, we performed RNA-Seq of UTEX 2797 cultures grown in six different conditions and four diurnal timepoints (Table S20). Starting 100 mL cultures were inoculated at 10,000 cells/mL and grown at 20 °C using a 12:12 light dark cycle and irradiance of 200 µmol photos $m^{-2}$ $s^{-1}$. Beginning five days post inoculation, cultures were maintained using semi-continuous replacement every three days by discarding 10% of the culture and replacing with fresh media. Cell densities were measured every three days to track culture growth. Upon reaching densities of ~$1x10^6$ cells/mL, cultures were harvested by centrifugation at 4500 x g for 5 minutes and snap freezing in liquid nitrogen. Total RNA was extracted from pelleted cells using the following protocol: https://dx.doi.org/10.17504/protocols.io.bv3hn8j6 (32). Stranded RNA-Seq libraries were constructed and sequenced by Novogene Corporation Inc. (Sacramento, CA) using a NEBNext Ultra TM RNA Library Prep Kit (NEB, USA) following manufacturer's recommendations. Libraires were sequenced on the Illumina NovaSeq 600 platform to produce 150 bp paired-end reads. RNA-Seq reads were aligned to the UTEX 2797 scaffolded assembly using STAR v2.7.8a (33) with the maximum intron length set to 10 kbp.

Gene model and protein prediction was first conducted on the UTEX 2797 scaffolded assembly with BRAKER2 v2.1.5 (34, 35). BRAKER2 was supplied the UTEX 2797 scaffolded assembly with repeats soft-masked, a custom protein database comprised of Swiss-Prot and all haptophyte predicted proteins from MMETSP (16), and the UTEX 2797 Illumina RNA-Seq data aligned to the soft-masked genome. All subsequent predictions for other strains via BRAKER2 utilized the resulting Augustus species-specific training configuration file (36) and the same custom protein databases.

**Characterization of Assembly Completeness.** Telomeric repeats were identified from scanning the final assemblies of 12B1 and UTEX 2797 with TRFFinder v4.09 (37) using the following parameters: 2 7 7 80 10 50 500 -f -d -m -h. Any tandem repeat within the first or last 2 kbp of a scaffold's length and whose repeat block was any permutation of the conserved Haptophyta telomere repeat TTAGGG (38) was selected. Positions of predicted telomeres are in Table S4. Conservation of core genes was performed using BUSCO v4.0.6 using the eukaryota_odb10 dataset (39, 40).

**Synteny.** Pairwise synteny between the Hi-C scaffolded genomes of UTEX 2797 and 12B1 was identified and visualized with the JCVI pipeline (41). Syntenic blocks within and between genome assemblies were identified with SynMap2 on the online Comparative Genomics Platform (CoGe) using the Relative Gene Order algorithm and Quota Align Merge with default settings to merge syntenic blocks (42). Synteny visualizations between UTEX 2797 haplotypes on syntenic scaffolds were performed with XMatchView (43).

**Phylogenomic analysis**. The phylogenetic workflow is summarized in Fig. S14. Orthologous sequences were identified using OrthoFinder v2.4.1 (44) with the single longest predicted coding sequence (CDS) per gene as input and using the blast_nucl sequence search option. Gene trees were constructed for every orthogroup containing sequences from at least 10 strains. Orthogroup CDS nucleotide sequences were aligned with MAFFT v7.471 (45) in the GUIDANCE v2.02 alignment software suite using the codon aware method (46). The length of the ungapped alignment was evaluated using TrimAL v1.4.rev15 (47), and orthogroups with ungapped alignments less than 150 bp were excluded from further analysis. Maximum likelihood (ML) gene phylogenies were constructed with IQ-TREE v1.6.12 (48) using the trimmed multiple sequence alignment as input. ModelFinder (49) was used to determine the best-fit nucleic acid substitution model, and 1000 replicates of both SH-aLRT and ultrafast bootstrapping analyses were performed. Gene trees were rooted based on the concatenation-based species tree (see below) using Notung v2.9.1.5 with its duplication, transfer, loss and ILS aware parsimony-based root optimization algorithm using default costs for all events (50, 51). In total, gene trees were built for 15,074 orthogroups that passed the strain count and alignment length filtering thresholds. The number of synonymous substitutions per synonymous site (*Ks*) was assessed for each pair of sequences within each orthogroup using the ungapped alignment as input. *Ks* was calculated according to LPB93 method using the yn00 module in PAML v4.9 (52–54).

The *P. parvum* species tree was constructed based on the combined signal from 2699 single-copy orthogroups (SCOGs) using both concatenation- and coalescence-based approaches. For the concatenation approach, the ML phylogeny was constructed in IQ-TREE v2.2.0 (55) based on a concatenated nucleotide data matrix consisting of 96 partitions and 2,982,918 sites with no missing data. The best partition model was selected using a relaxed hierarchical clustering algorithm as implemented by ModelFinder in IQ-TREE v2.2.0. The concatenation-based species tree was rooted on the branch with the highest rootstrap support as calculated in IQ-TREE v2.2.0 using the same best partition scheme as above but with a linked non-reversible DNA substitution model 12.12 across all partitions. The --root-test option was enabled to perform a tree topology test and compare the log-likelihoods of the trees based on every possible root location (56). The coalescence-based phylogeny estimation was conducted using ASTRAL v5.7.1 (57), and the resulting tree was manually rooted to match the root placement of the concatenation-based approach. Gene and site concordance factors were calculated for both species trees with IQ-TREE v2.2.0 using the 2699 single-copy genes used to construct the species trees.

Multi-labeled (MUL) species trees were built using GRAMPA v1.3 with UTEX 2797 as the h1 polyploid clade (58). Two MUL tree analyses were run using different sets of Notung rooted gene trees as input: SCOGs only (2699 trees) and all orthogroups (15,040 trees). The MUL tree with the best parsimony score was the same using either gene tree set (Table S18). Sister taxa of each UTEX 2797 gene were identified using the Bio.Phylo Biopython toolkit (59) (see Data Availability; Fig. S18). The proportion of genes along each UTEX 2797 scaffold that grouped with subclade A1 or A2 were calculated with BEDTools intersect (60) using 500 kbp windows and requiring a 50% minimum overlap for each gene. Subclade proportions and scaffold synteny were visualized using Circos v0.69-9 (61).

**Breadth of coverage.** Breadth of coverage was calculated from alignments of the filtered Illumina reads to the 12B1 and UTEX 2797 scaffolded assemblies. To control for library differences between strains, the filtered Illumina reads were randomly subsampled to equal read counts (N = 44 million pairs) using reformat.sh, a tool of the BBMap software suite v38.87, with sample seed set to 13 (18). Alignments were generated using BWA-MEM v0.7.15 (12). Depth of coverage was calculated on a per-base level for both 12B1 and UTEX 2797 assemblies using samtools. Breadth of coverage was calculated as the proportion of base-pairs in the final assembly that had coverage greater than N coverage. BOC using coverage N = 0, 5, 10, 20, and 50 were all evaluated. Final breadth of coverage tables for each assembly can be found in Tables S11.

**Heterozygosity and read coverage analyses**. Heterozygosity and coverage of maximal unique k-mers were estimated using the subsampled, filtered Illumina gDNA reads (see Characterization of Assembly Completeness Methods section) using KMC v3.1.1 (62) with a k-mer length (-k) of 21, minimal k-mer occurrence (-ci) of 1, and maximal k-mer occurrence (-cs) of 10,000.
`

Read coverage of single copy orthogroups (SCOGs) was calculated by first aligning the subsampled, filtered Illumina gDNA reads to each strain's assembly using BWA-MEM v0.7.15 (12). Mean read depths were calculated for each SCOG with BEDTools intersect (60) with the -mean option enabled. The median read depth of the 2699 SCOGs was then used as an estimate of the average coverage of the genome. The haploid genome size (G) was calculated using the Lander-Waterman equation (63): $G = LN/C$ where LN is the total combined length (bp) of the input Illumina reads and C is coverage as estimated by SCOG read depth.

**Cell size estimation.** Cell sizes were measured when cultures were in early stationary phase, and cultures were kept in each strain's standard growth medium (Table S8). Samples were drawn from cultures after gentle swirling, stained with Lugol's acidic solution in a 1.5 mL microcentrifuge tube, vortexed briefly, and promptly loaded into a glass slide with cover slip for immediate imaging. Final size data were collected from ≥ 15 images for each strain (50 cells total). Images covering non-overlapping, haphazardly chosen areas of the slide were pre-processed to remove artifacts and converted to binary using an automated threshold selection process in ImageJ2 (64). Radii of major and minor axes of each cell were used to estimate cell volumes as ellipsoids using the formula: $V = \frac{4}{3}\pi(r_{minor}{}^{2})(r_{major})$.

**Functional annotation and enrichment tests.** Gene functional annotations were assigned via InterProScan v5.50-84.0 (65) using default settings and KofamScan (66) with the threshold-scale set to 0.9. Tests for enrichment of higher-level functional categories were performed using the core go.obo ontology (Gene Ontology Consortium) and the KEGG PATHWAY metabolic hierarchy downloaded via the KEGG API. Hypergeometric tests were performed in python using the SciPy library hypergeom, and p-values were adjusted for multiple comparisons using the StatsModels library multitest with the Benjamini & Hochberg (BH) method (67).

**Horizontal gene transfer.** We assessed the genomes for possible HGT events using the Alien Index (AI) score as previously described (68). Briefly, each predicted protein sequence was queried against the same custom protein database used for BlobTools (see above) with DIAMOND (v2.0.8.146) (14). A custom python script sorted the DIAMOND results based on the normalized bitscore (*nbs*), where *nbs* was calculated as the bitscore of the single best scoring HSP to the subject sequence divided by the best bitscore possible for the query sequence (i.e., the bitscore of the query aligned to itself). The AI score is given by the formula: $AI = nbsO - nbsH$, where *nbsO* is the normalized bit score of the best hit to a species outside of the Haptista lineage (NCBI:txid2608109), *nbsH* is the normalized bit score of the best hit to a species within the Haptista lineage skipping all hits to the *Prymnesium* genus (NCBI:txid35143). AI scores range from -1 to 1, being greater than 0 if the predicted protein sequence had a better hit to a non-Haptista sequence, suggestive of either horizontal gene transfer (HGT) or contamination (68). Because donor and recipient lineages cannot be differentiated based on AI score alone, and given the relatively common occurrence of HGT from haptophytes to dinoflagellates (particularly dinoflagellates containing haptophyte-derived chloroplasts) (69), we also skipped all hits to the Dinophyceae lineage (NCBI:txid 2864) when calculating AI scores. We filtered our HGT candidates (AI > 0) to those that were most likely to be phylogenetically informative by requiring AI > 0.1 and total hits ≥ 50. In addition, if the top hit was to a eukaryote, we further required than ≥ 5 haptophytes from outside the *Prymnesium* genus be present in the top 200 hits. This extra requirement allowed us to evaluate whether haptophytes formed a monophyletic clade or whether *Prymnesium* grouped separately, which would provide stronger support for HGT (70). See Data Availability for access to the database and scripts.

Phylogenetic trees of protein sequences were constructed for all filtered AI-flagged HGT candidates. Full-length proteins corresponding to the top 200 hits (E-value < $1 \times 10^{-10}$) to each query sequence were extracted from the local database using esl-sfetch (71). Protein queries with less than 50 significant hits were skipped. Protein sequences were aligned with MAFFT v7.471 using the E-INS-i strategy and the BLOSUM30 amino acid scoring matrix (45) and trimmed with trimAL v1.4.rev15 using its gappyout strategy (47). Proteins with trimmed alignments < 150 amino acids in length were excluded. The topologies of the remaining genes were inferred using maximum likelihood as implemented in IQ-TREE v1.6.12 (48) using an empirically determined substitution model and 1000 rapid bootstrap replications. The phylogenies were midpoint rooted and branches with local support < 95 were collapsed using the ape and phangorn R packages (72, 73). Phylogenies were visualized using ITOL version 4 (74) and inspected manually to identify phylogenetically supported HGT candidate proteins.

References cited in SI figure legends (75) (76) (77).

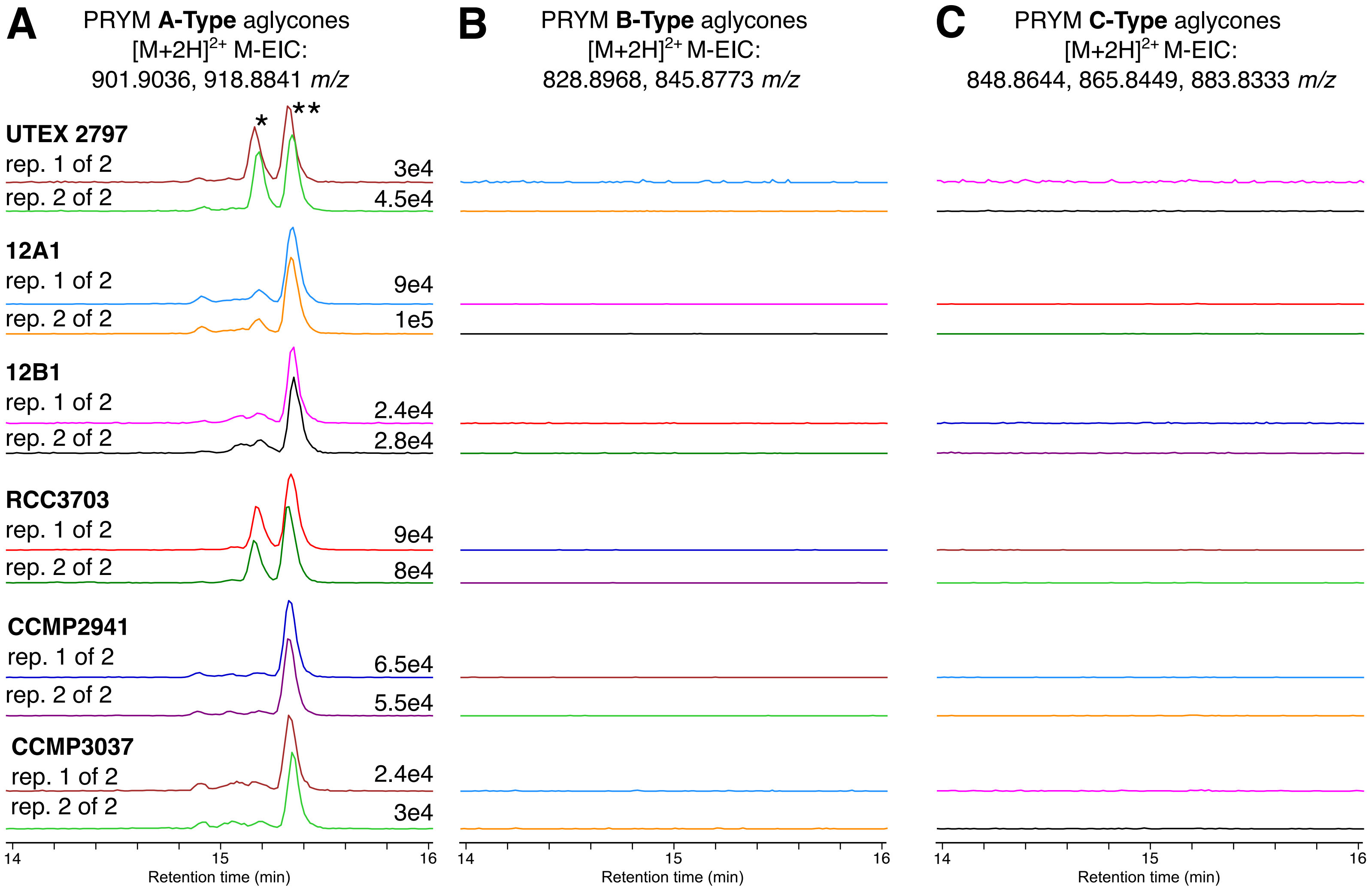


**Fig. S1. Chemotyping of *P. parvum* strains.** All strains analyzed (UTEX 2797, 12A1, 12B1, RCC3703, CCMP294, CCMP3037) were found to produce A-type prymnesins (PRYM), via LC-HRAM-MS multi-extracted-ion-chromatograms (M-EICs) of representative $[M+2H]^{2+}$ aglycone in-source fragments ions. M-EICs are the sum of 2 or more EICs (A) M-EICs of 2 calculated *m/z* values for representative PRYM A-type aglycone $[M+2H]^{2+}$ ions ($Cl_2$: 901.9036 *m/z*, $Cl_3$: 918.8841 *m/z;* from SI ref 75). Some strain-to-strain variation in PRYM A-type congener proportions is present. *=largely due to 901.9036 *m/z* **=largely due to 918.8841 *m/z* (B) E-MICs of 2 representative PRYM B-type aglycone $[M+2H]^{2+}$ ions ($Cl_2$: 828.8968 *m/z*, $Cl_3$: 845.8773 *m/z;* from SI ref 75). (C) E-MICs from 3 representative PRYM C-type aglycone $[M+2H]^{2+}$ ions ($Cl_1$: 848.8644 *m/z*, $Cl_2$: 865.8449 m/z, $Cl_3$: 883.8333 m/z; from SI ref 75). Norm=normalization. Rep=Biological replicate.

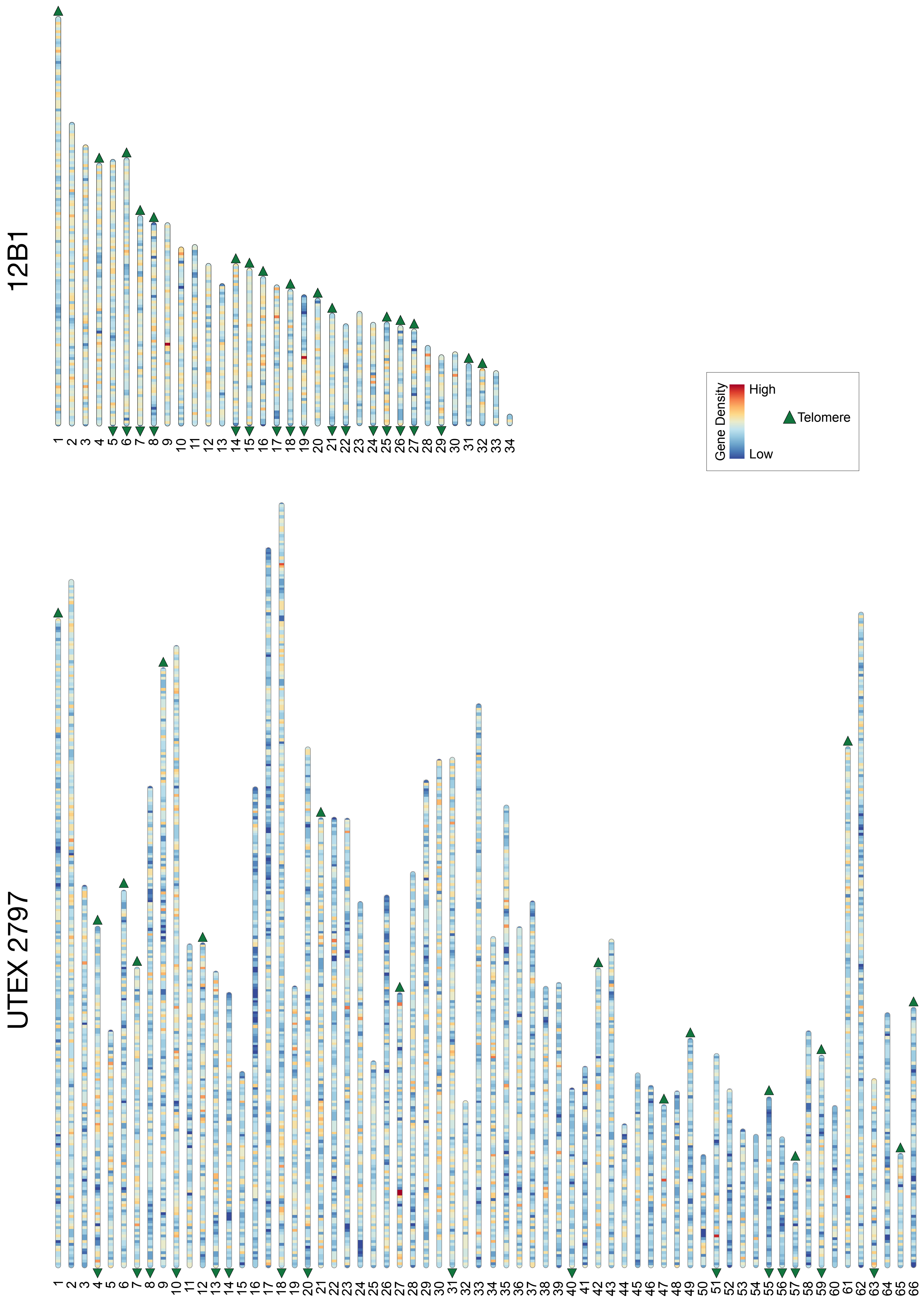


**Fig. S2. *Prymnesium parvum* genome assembly karyotypes for strains 12B1 and UTEX 2797.** Karyotypes depict gene density and presence/absence of telomere repeats (triangles) at scaffold termini.

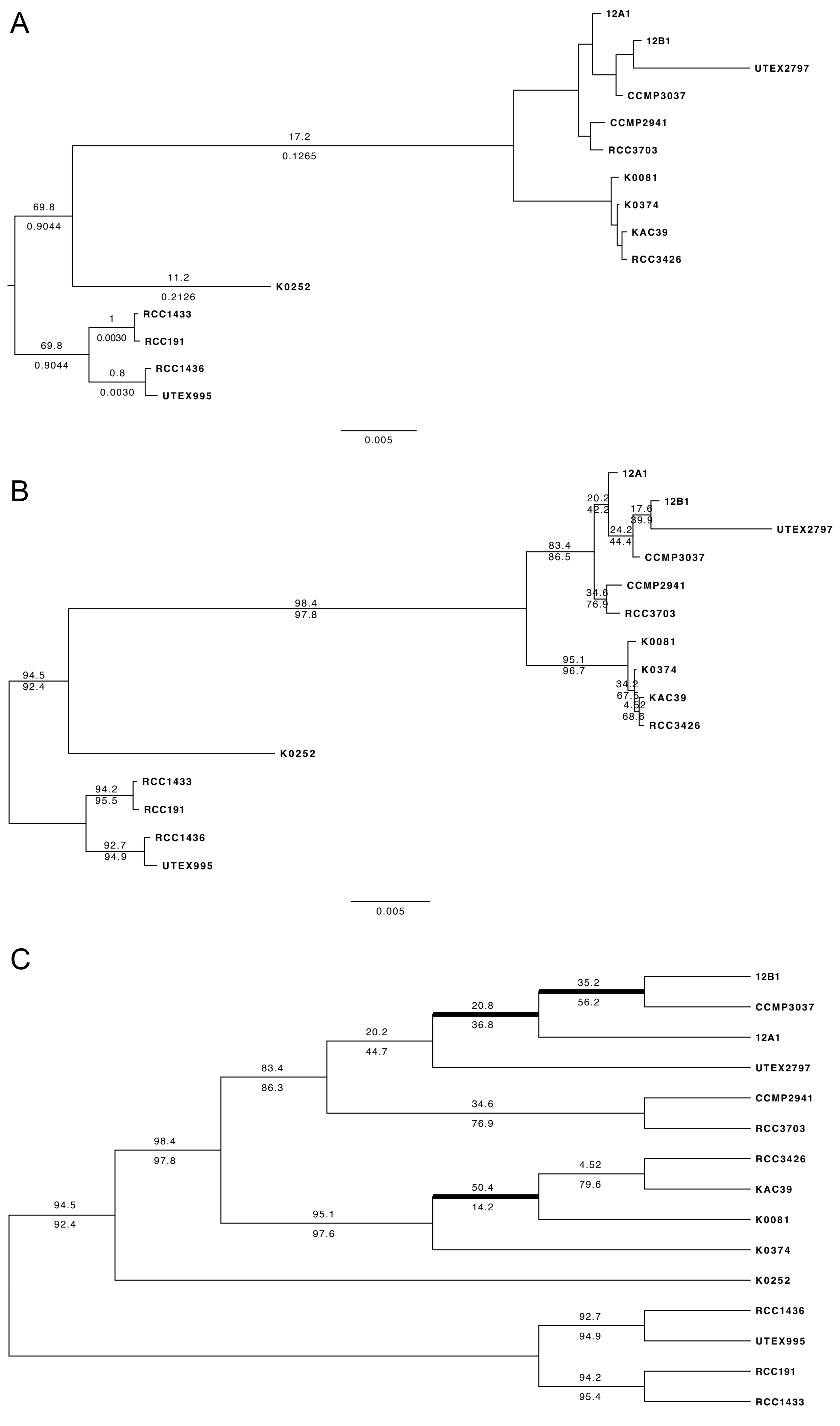


**Fig. S3. Phylogenetic relationships of 15 *Prymnesium parvum* strains based on 2699 single-copy genes.** A) ML phylogeny reconstructed based on the concatenated nucleotide data matrix. Numbers above branches indicate IQ-TREE rootstrap support, and numbers below branches indicate p-value of approximately unbiased (AU) test for alternative root locations (branches with p-values < 0.05 were considered a significantly worse root location compared to displayed root). Unlabeled branches had rootstrap support = 0 and pAU < 0.001. B) Same ML phylogeny as shown in part A; numbers above and below branches indicate the gene concordance factor and site concordance factor, respectively. C) Species tree inferred from an ASTRAL coalescence-based analysis; numbers above and below branches indicate the gene concordance factor and site concordance factor, respectively. Thick branches show conflicts between the concatenation-based phylogeny.

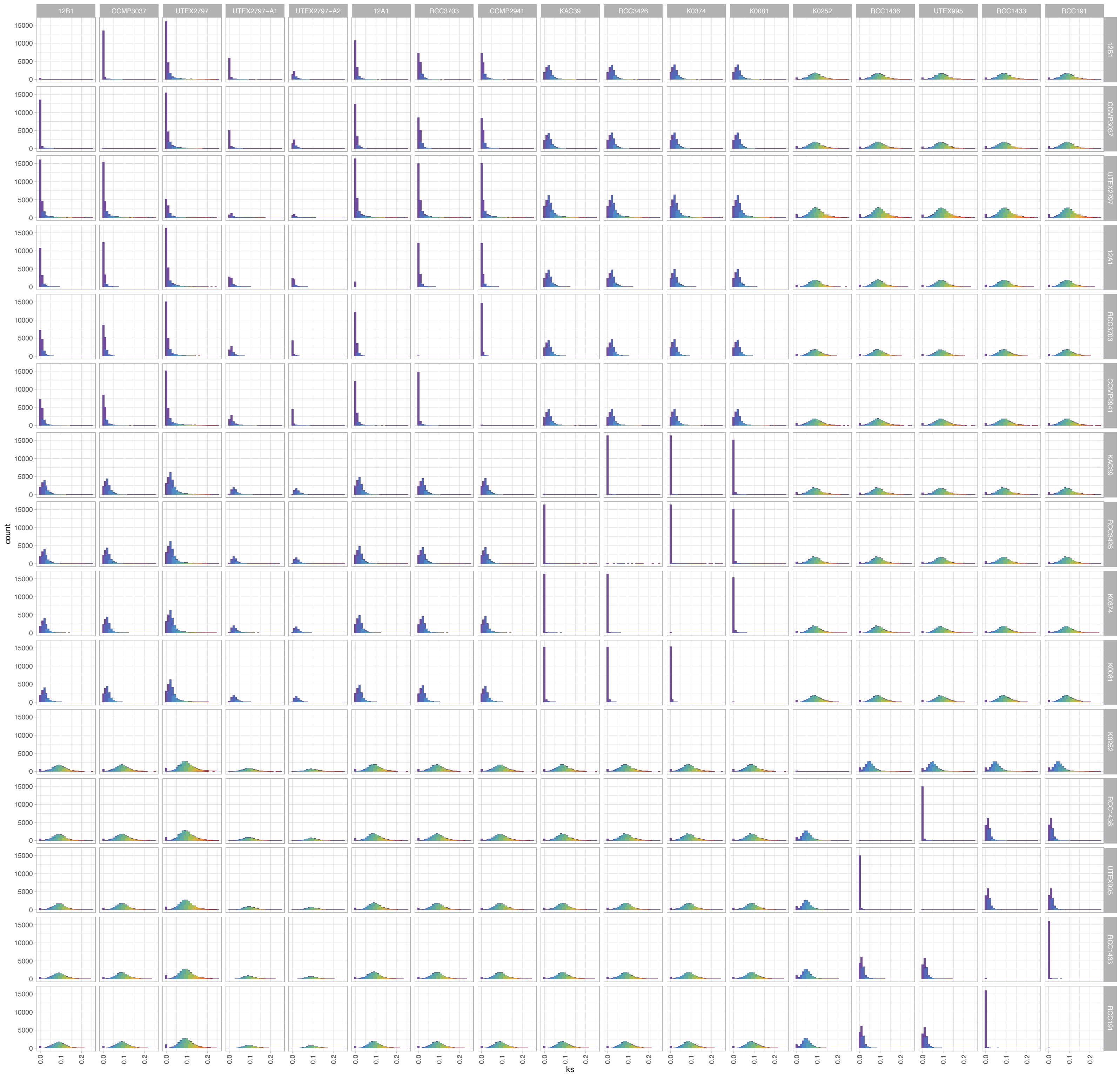

**Fig. S4. *Ks* plots.** Distribution of substitutions per synonymous site (*Ks*) between all strains. To avoid signal from distant paralogs and/or poor alignments, gene pairs with *Ks* values > 0.25 were excluded.

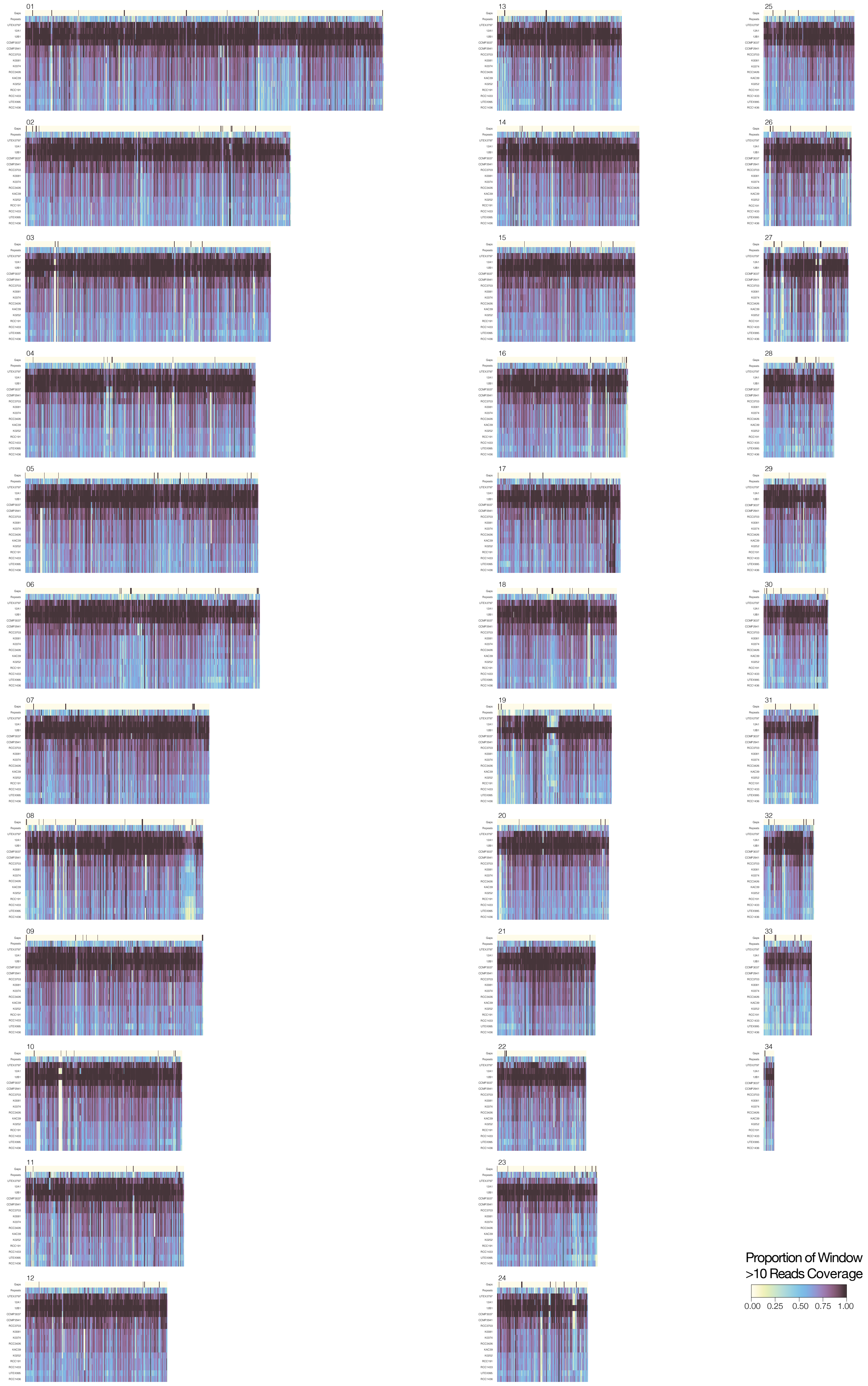


**Fig. S5. Breadth of coverage map of reference strain 12B1.** Illumina coverage for all strains along all 12B1 scaffolds. The color scale reflects the percentage of bases in 10 kbp windows that have at least 10x depth. Black boxes in the first row represent the location of assembly gaps, and the second row indicates repeat density (percentage of bases per window that are repetitive). All other rows represent breadth of coverage for each strain.

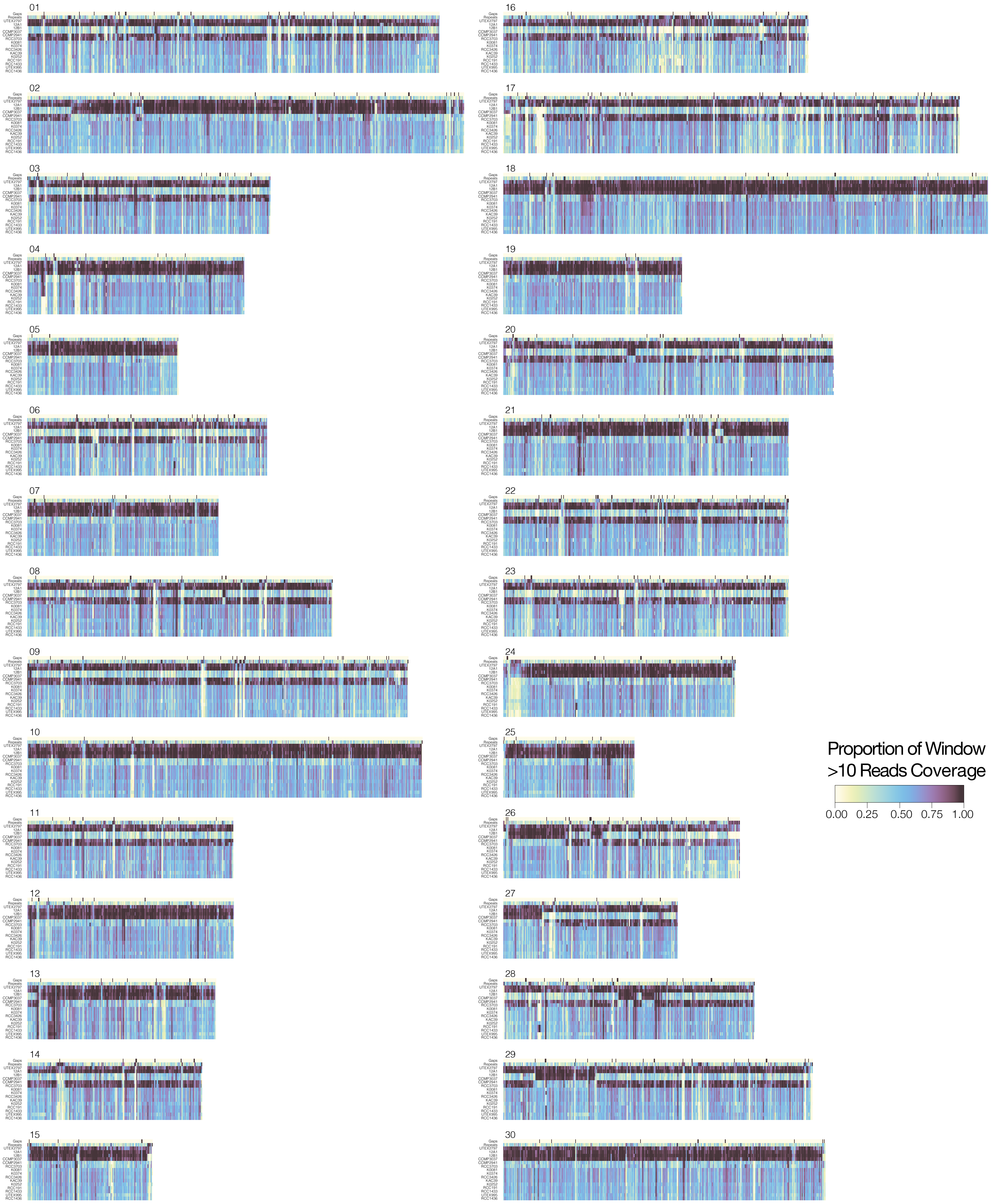


**Fig. S6. Breadth of coverage map of reference strain UTEX 2797.** Illumina coverage for all strains along all UTEX 2797 scaffolds. The color scale reflects the percentage of bases in 10 kbp windows that have at least 10x depth. Black boxes in the first row represent the location of assembly gaps, and the second row indicates repeat density (percentage of bases per window that are repetitive). All other rows represent breadth of coverage for each strain.

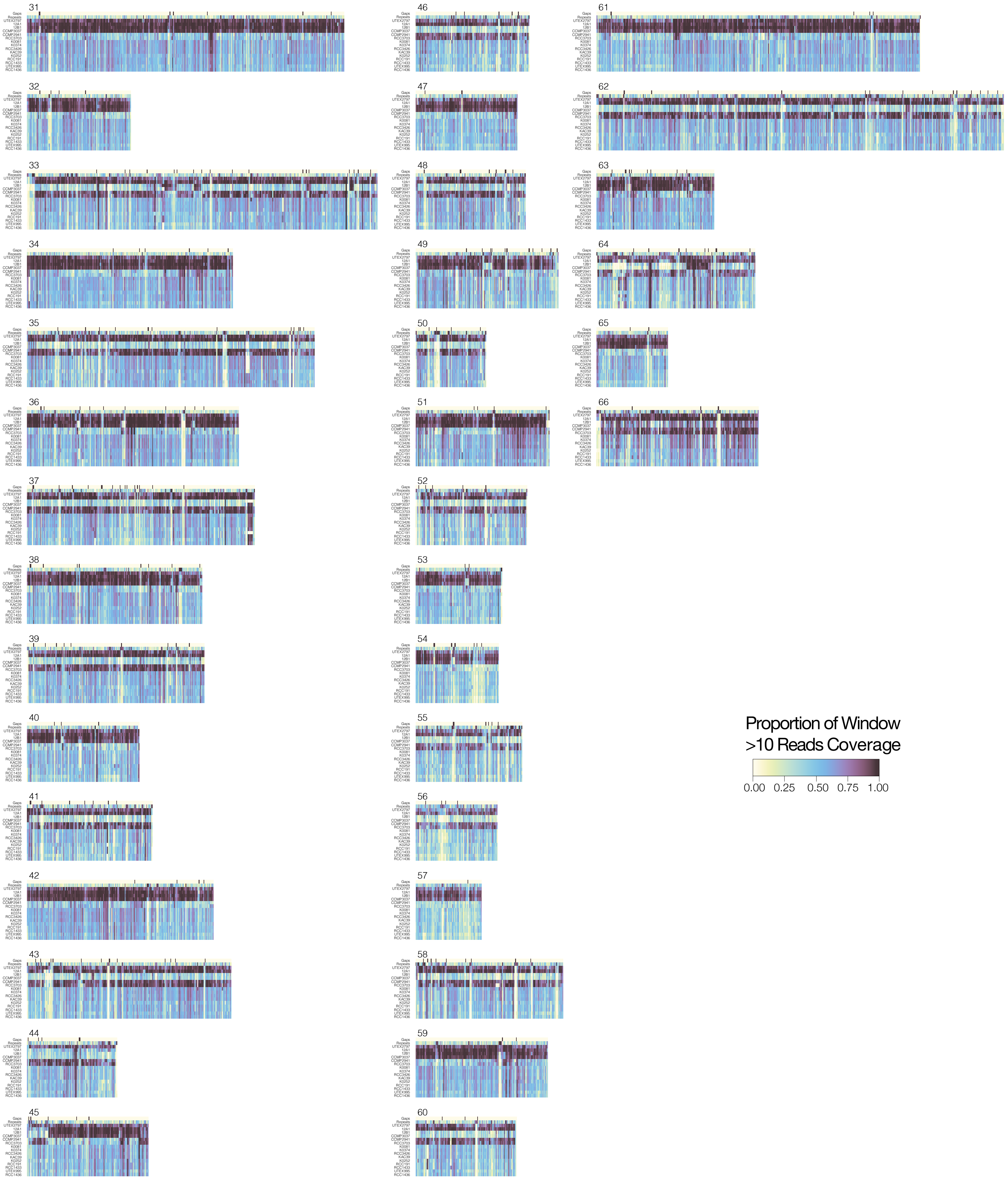

Fig. S6 cont. Breadth of coverage map of reference strain UTEX 2797. Illumina coverage for all strains along all UTEX 2797 scaffolds. The color scale reflects the percentage of bases in 10 kbp windows that have at least 10x depth. Black boxes in the first row represent the location of assembly gaps, and the second row indicates repeat density (percentage of bases per window that are repetitive). All other rows represent breadth of coverage for each strain.

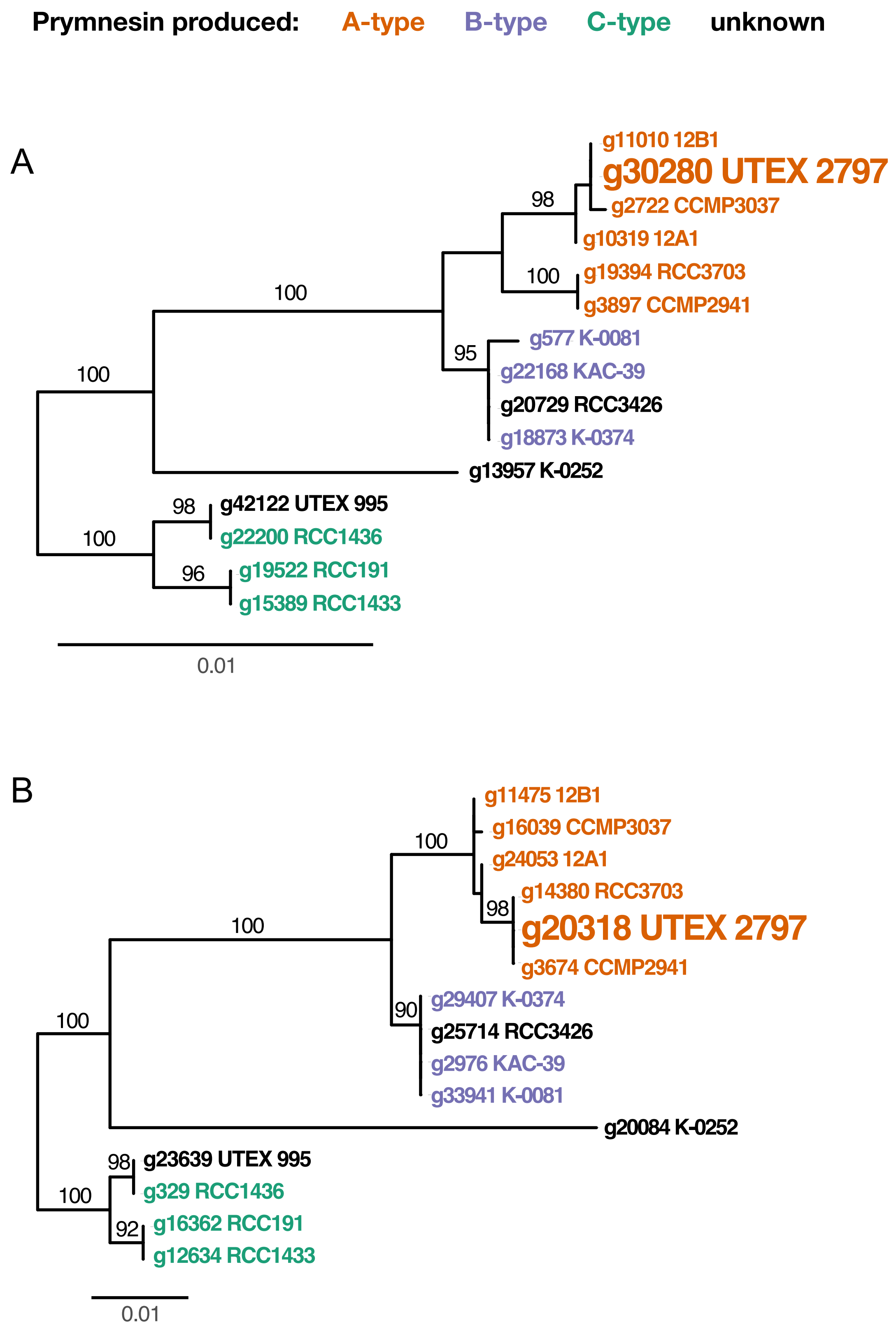


**Fig. S7. Example gene trees depicting variable location of UTEX 2797.** A) UTEX 2797 groups with 12B1, CCMP 3037 and 12A1 in gene tree for orthogroup OG0017620. B) UTEX 2797 groups with RCC3703 and CCMP2941 in OG0015960. Numbers above branches indicate ultrafast bootstrap support >90 for the descendent node.

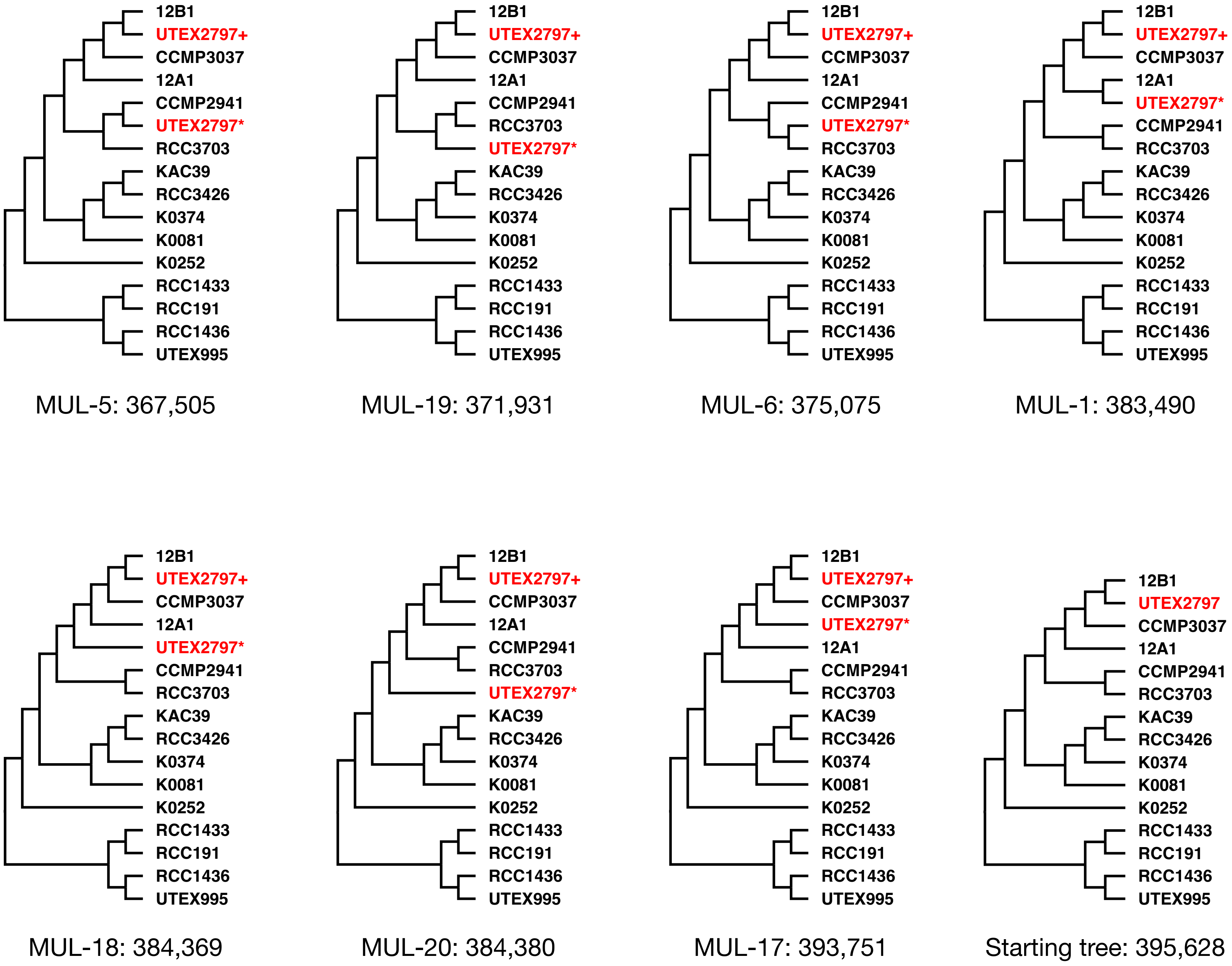


**Fig. S8. Multi-labeled (MUL) species trees.** Tree on the bottom right is the single-labeled species tree (ST). All other trees are MUL trees in which UTEX 2797 appears twice (red). The parsimony score is included at the bottom of each tree and indicates the total duplication and loss cost to reconcile the input gene trees to each species tree (smaller values indicates more parsimonious solutions). In this analysis all 15,035 gene trees served as input; a second analysis using only 2699 single copy orthogroup trees showed a similar pattern (see Table S12 for all possible MUL tree scores).

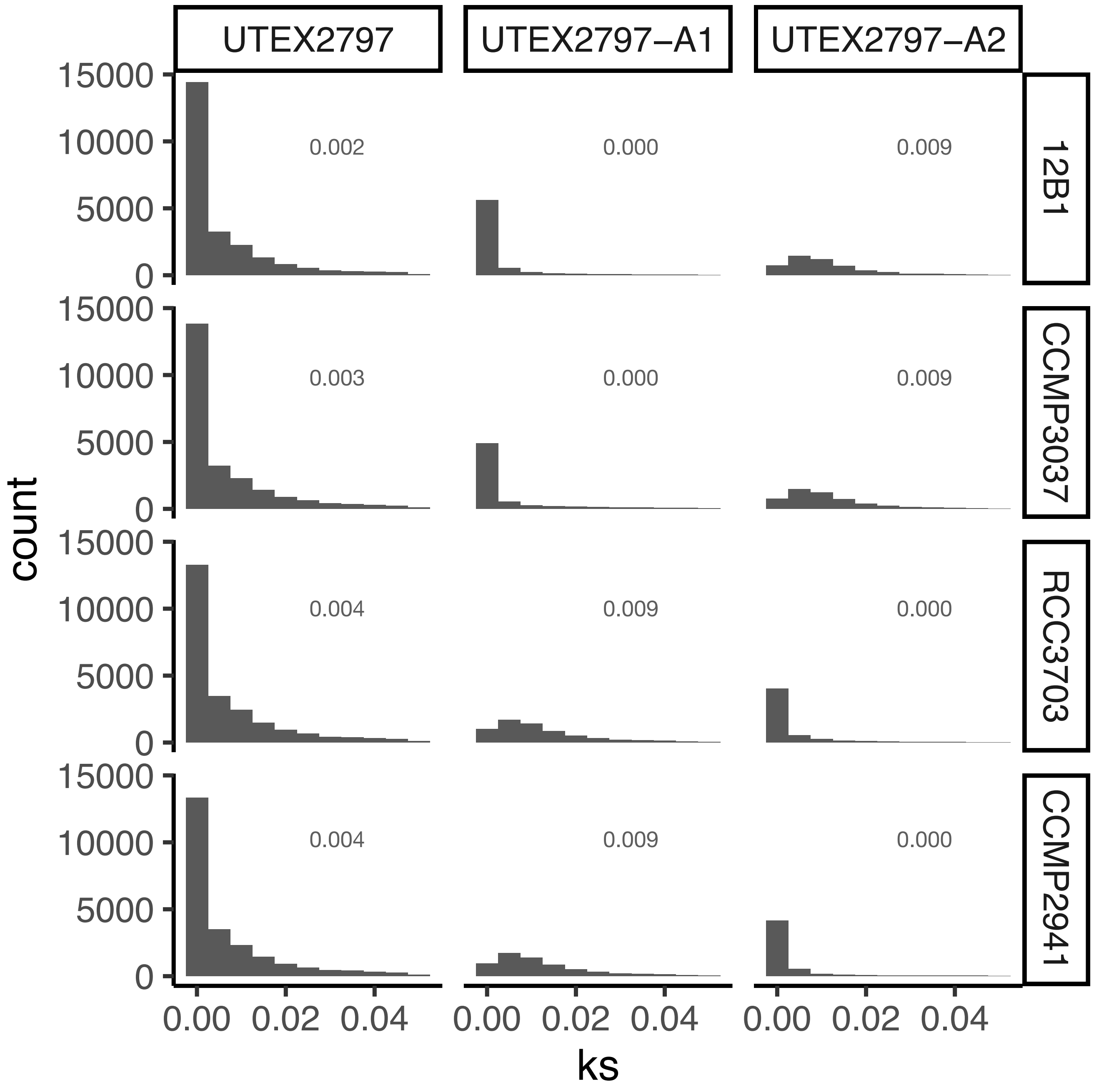


**Fig. S9. *Ks* plots for UTEX 2797 subgenomes.** Distribution of substitutions per synonymous site (*Ks*) between all UTEX 2797 genes compared to those that group with A1 clade strains and those that group with A2 type strains. No branch support threshold was imposed for A1/A2 clade calls. These gene sets were aligned to homologs in A1 strains (12B1 and CCMP3037) and A2 strains (RCC3703 and CCMP2941). To avoid signal from distant paralogs and/or poor alignments, gene pairs with *Ks* values > 0.05 were excluded. Gray numbers indicate median *Ks* values for each distribution.

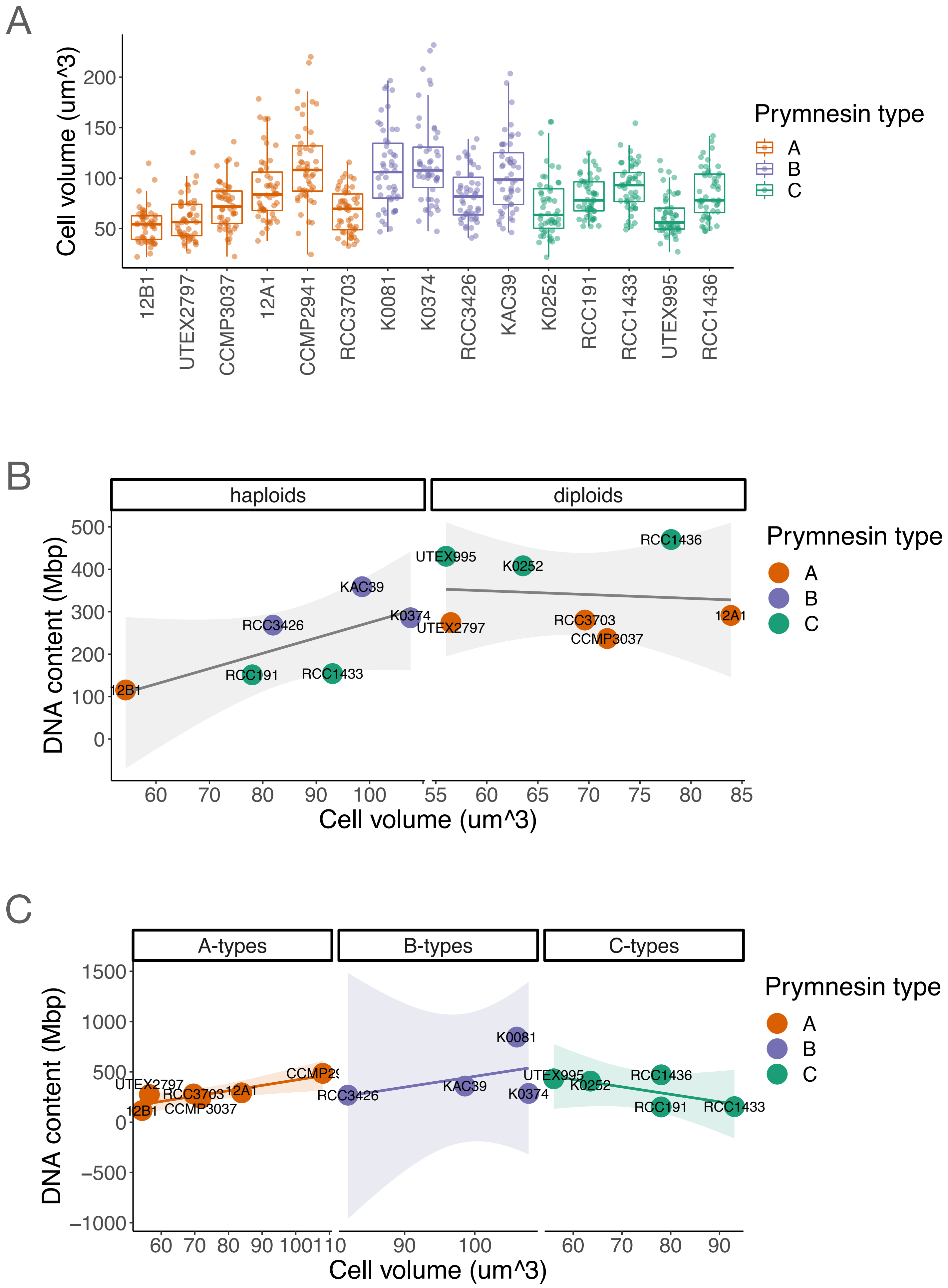


**Fig. S10. Variation in *Prymnesium parvum* cell volume and genome size.** A) Cell volume distributions per strain. B) The relationship between cell volume and total DNA content plotted as a factor of predicted ploidy. C) The relationship between cell volume and total DNA content plotted as a factor of prymnesin type. Lines of best fit and associated confidence intervals in parts B and C were plotted using a linear model using the geom_smooth() function in ggplot2.

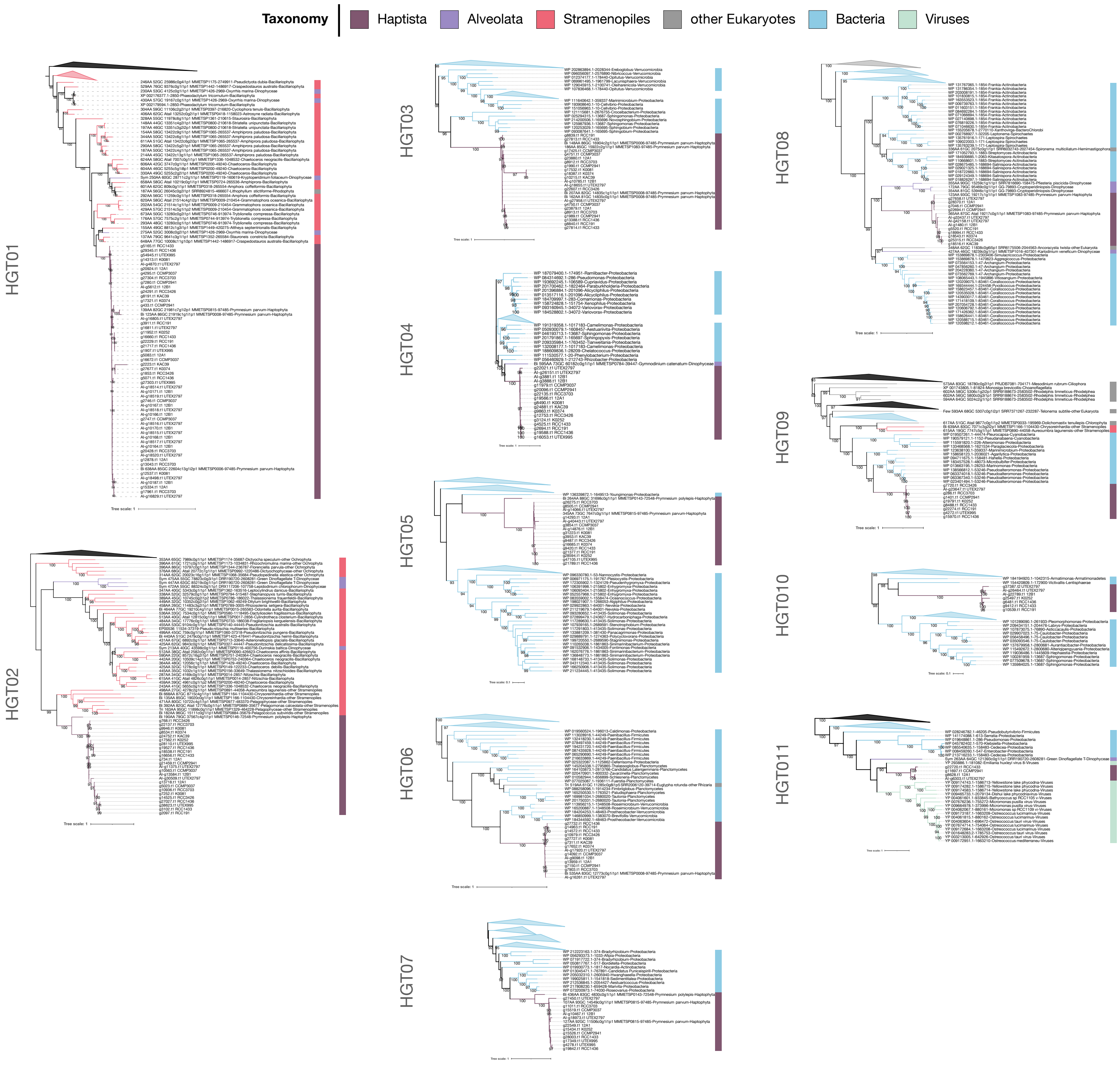


**Fig. S11. Maximum likelihood phylogenies of horizontal gene transfer events in *Prymnesium parvum*.** Nodes with IQ-TREE support values > 90 are indicated by numbers on the preceding branch. The branches and outer color bars are color-coded to match the taxonomic classification of each sequence. Trees are midpoint rooted. Triangles indicate collapsed clades that have been reduced for clarity. Triangle color indicates the taxonomy of the collapsed taxa; black triangles indicate that downstream taxa belong to multiple lineages.

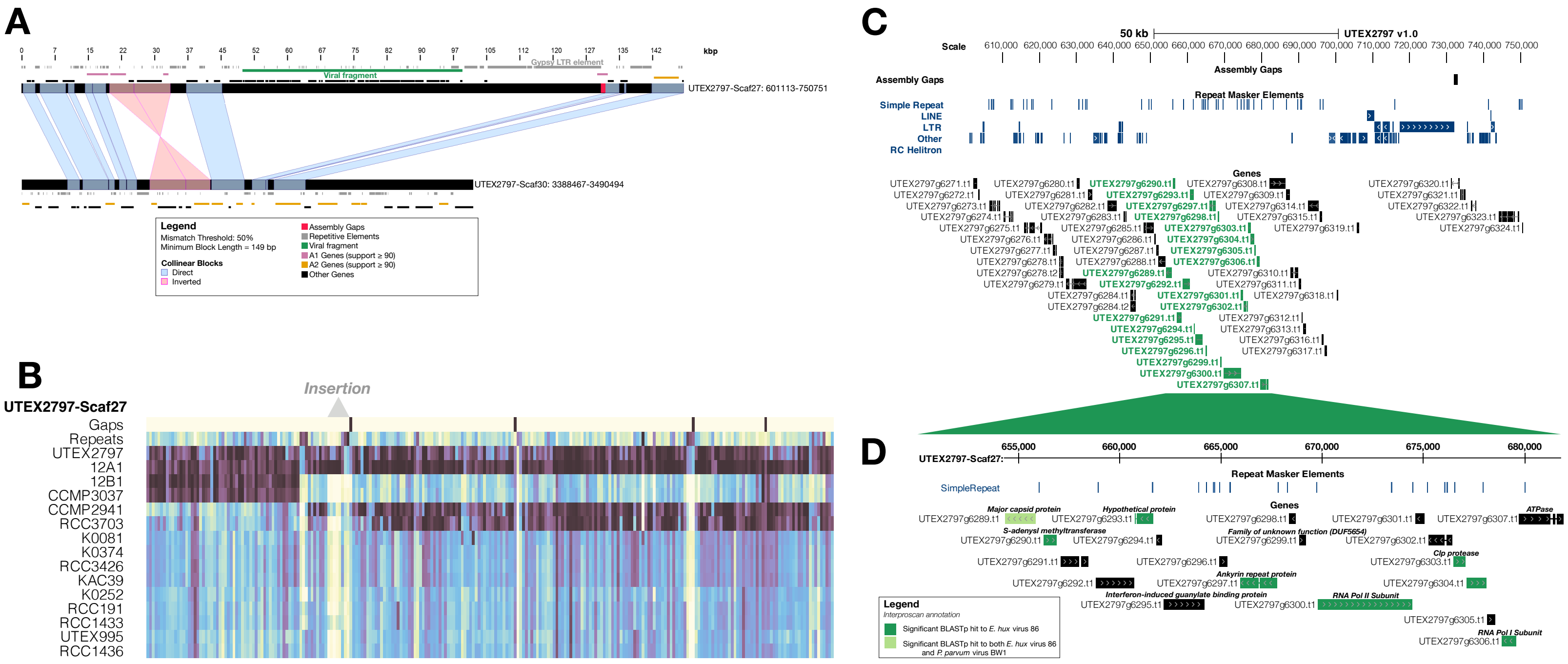


**E**

| query | Best hit to Ehux virus 86 | Bitscore (Ehux virus 86) | Best hit to Pparvum virus BW1 | Bitscore (Pparvum virus BW2) |
|---|---|---|---|---|
| UTEX2797g6289 | YP_293839.2 | 2.71E-136 | AQV04380.1 | 2.71E-47 |
| UTEX2797g6290 | YP_293810.1 | 1.44E-35 | -- | -- |
| UTEX2797g6293 | YP_293830.1 | 6.62E-15 | -- | -- |
| UTEX2797g6297 | YP_294212.1 | 0.00000104 | -- | -- |
| UTEX2797g6300 | YP_294192.1 | 0 | -- | -- |
| UTEX2797g6303 | YP 293886.1 | 6.49E-44 | -- | -- |
| UTEX2797g6306 | YP_294157.1 | 8.67E-17 | -- | -- |

**Fig. S12. Characterization of the HGT11 viral insertion.** (A) Synteny visualization of the HGT11 viral insertion on UTEX2797-Scaf27 (top) with the homeologous region on UTEX2797-Scaf30 (bottom) via XMatchView. Collinear sequences are joined by blue bands if in the same orientation or pink bands if inverted. Repetitive elements are indicated as gray boxes. Genes are differentially colored to indicate whether they group phylogenetically with A1 or A2 strains (pink or orange, respectively) or neither (black). The up- and downstream boundaries of the putative viral insertion is marked by a green bar, which reflect the first and last gene that has BLAST hits to viral genes. (B) Illumina coverage along UTEX 2797 surrounding the insertion (gray triangle) for all strains. Colors reflect the percentage of bases in 10 kbp windows that have at least 10x read depth. Black boxes in the first row represent the location of assembly gaps, and the second row indicates repeat density (percentage of bases per window that are repetitive). (C) Genomic view along UTEX2797-Scaf27 surrounding the insertion. Gaps (black) and repeats (blue) are denoted as separate tracks. Genes that are putatively of viral origin are green, while genes that are not are black. (D) Zoomed in view of the viral-like genes above. If the gene had a significant BLASTp hit to the *E. huxleyi* virus 86 (NC_007346.1) are dark green, while genes that had a significant hit to both *E. huxleyi* virus 86 and *P. parvum* virus BW1 (AQV04380.1) are light green. If a gene was functionally annotated through InterProScan, the annotation is provided above the gene model. (E) Table of best BLASTp hits to the *E. huxleyi* virus 86 and *P. parvum* virus BW1.

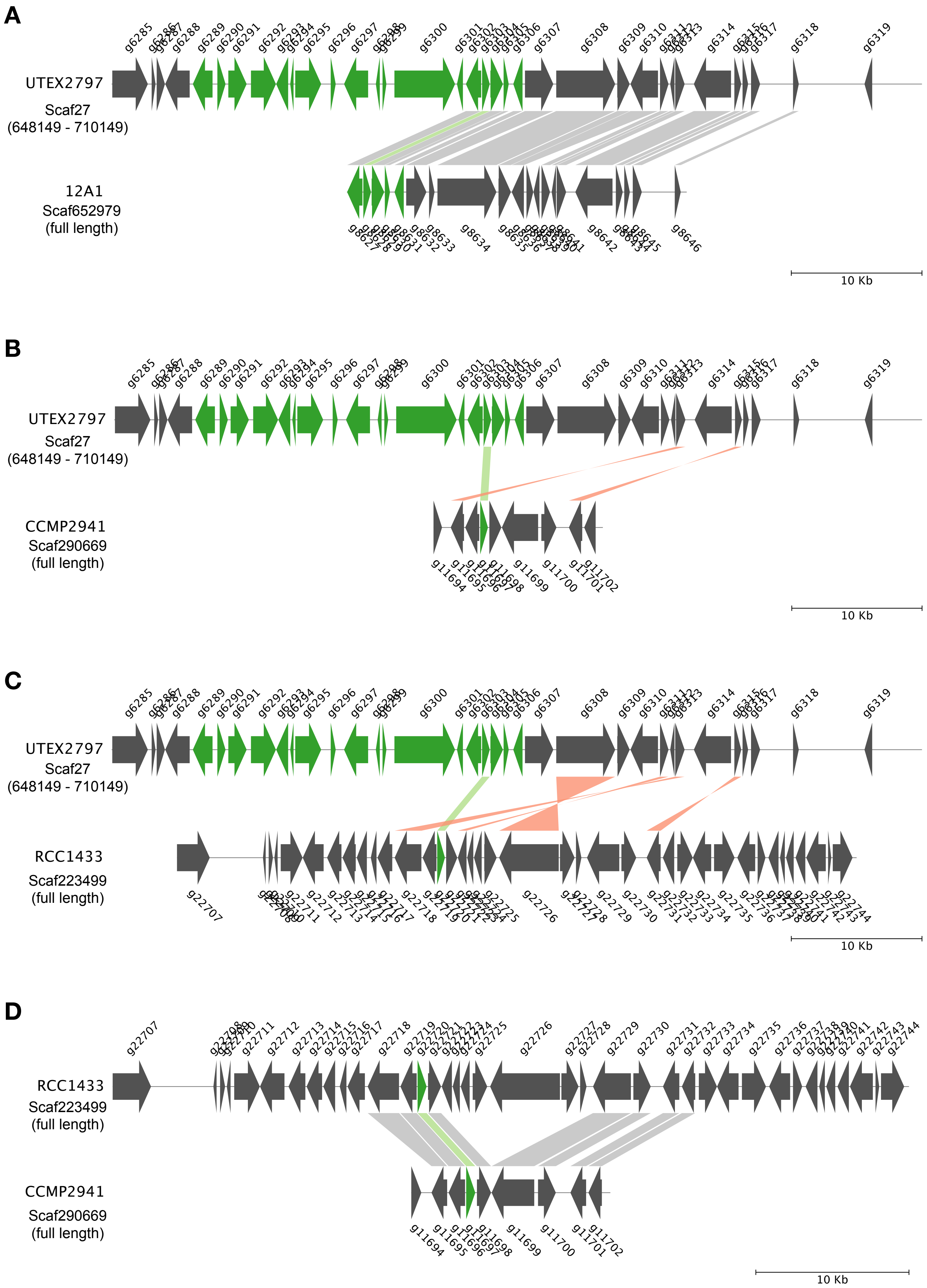


**Fig. S13. Comparison of shared synteny in region of HGT11 viral insertion.** Genome regions are visualized with pyGenomeViz (SI ref 76). HGT11 consists of g6303 in UTEX 2797; g8628 in 12A1; g11697 in CCMP2941; and g22720 in RCC1433 and is absent in all other eleven strains in the analysis (See Fig. S11). Light green bands indicate the location of HGT11 genes; all other collinear sequences are joined by grey bands if in the same orientation or orange bands if inverted. Green genes in UTEX 2797 indicate the genes putatively acquired via a single HGT event (See Fig. S12). In 12A1, Scaf652979 is syntenic across its entire length with UTEX 2797, including five genes of the putative viral insertion (part A), suggesting that 12A1 and UTEX 2797 likely share the same insertion event. In CCMP2941, Scaf290669 is syntenic across its entire length with Scaf223499 in RCC 1433, but neither regions are syntentic with UTEX 2797 despite sharing a common viral gene (parts B-D). This suggests that the HGT event giving rise to the viral gene in RCC1433 and CCMP2941 is likely separate from that of UTEX2797 and 12A1. However, long-read assemblies are required for 12A1, CCMP2941, and RCC1433 to rule out viral contamination in Illumina-only assemblies.

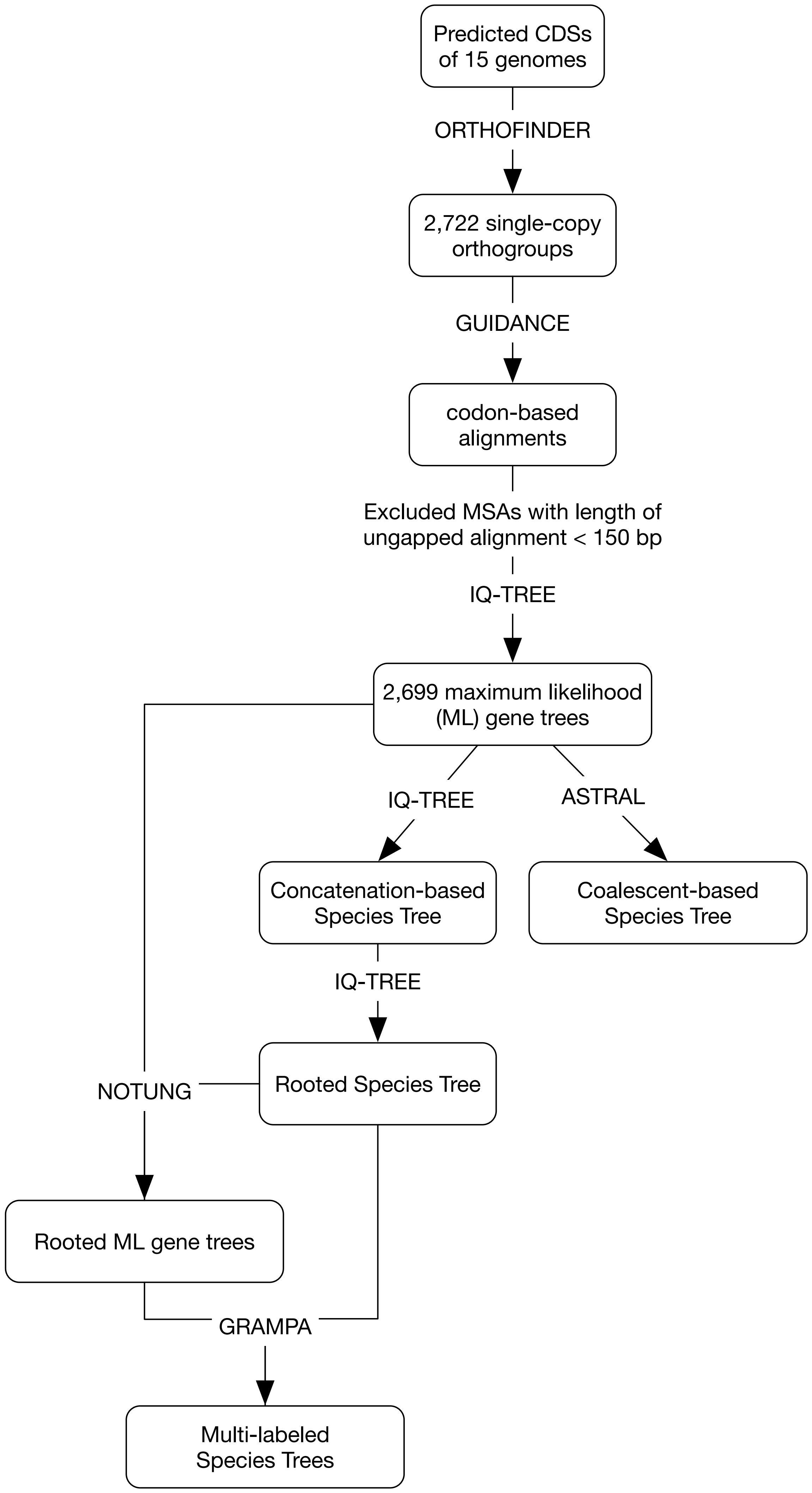


**Fig. S14. Phylogenetic workflow.**

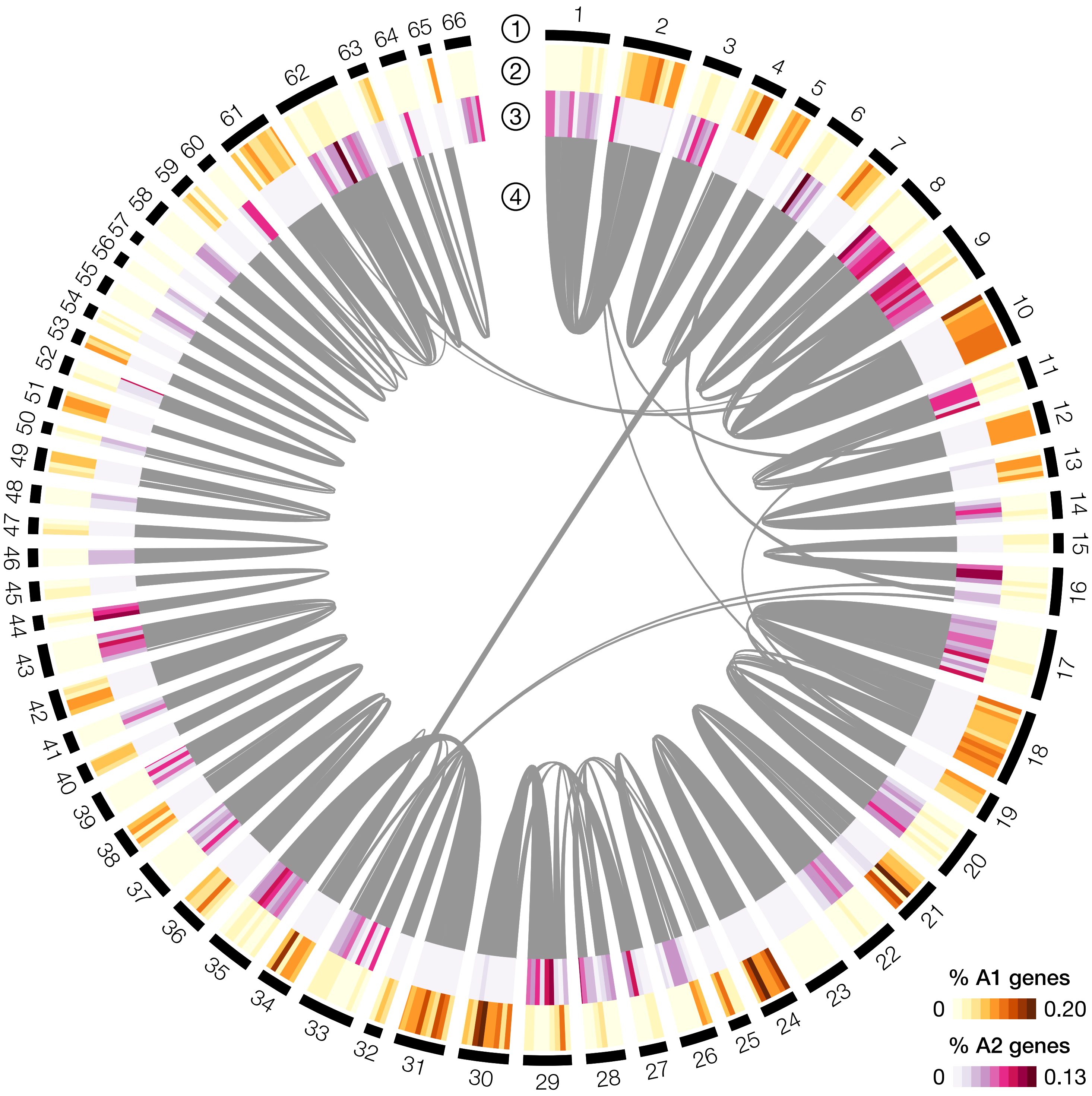


**Fig. S15. Circos plot imposing 90 branch support for A1/A2 calls.** Circos plot showing the 66 scaffolds of UTEX 2797 with four tracks (1) outer black track indicates scaffolds, (2) orange heatmap indicating the percentage of genes in 50 kbp windows that group within the A1 clade, (3) pink heatmap indicating the percentage of genes in the same 50 kbp windows that group within the A2 clade, and (4) grey bands indicate syntenic blocks (≥ 15 syntenic genes per block). Branch support threshold >=90 was required to assign genes to A1/A2 clades.

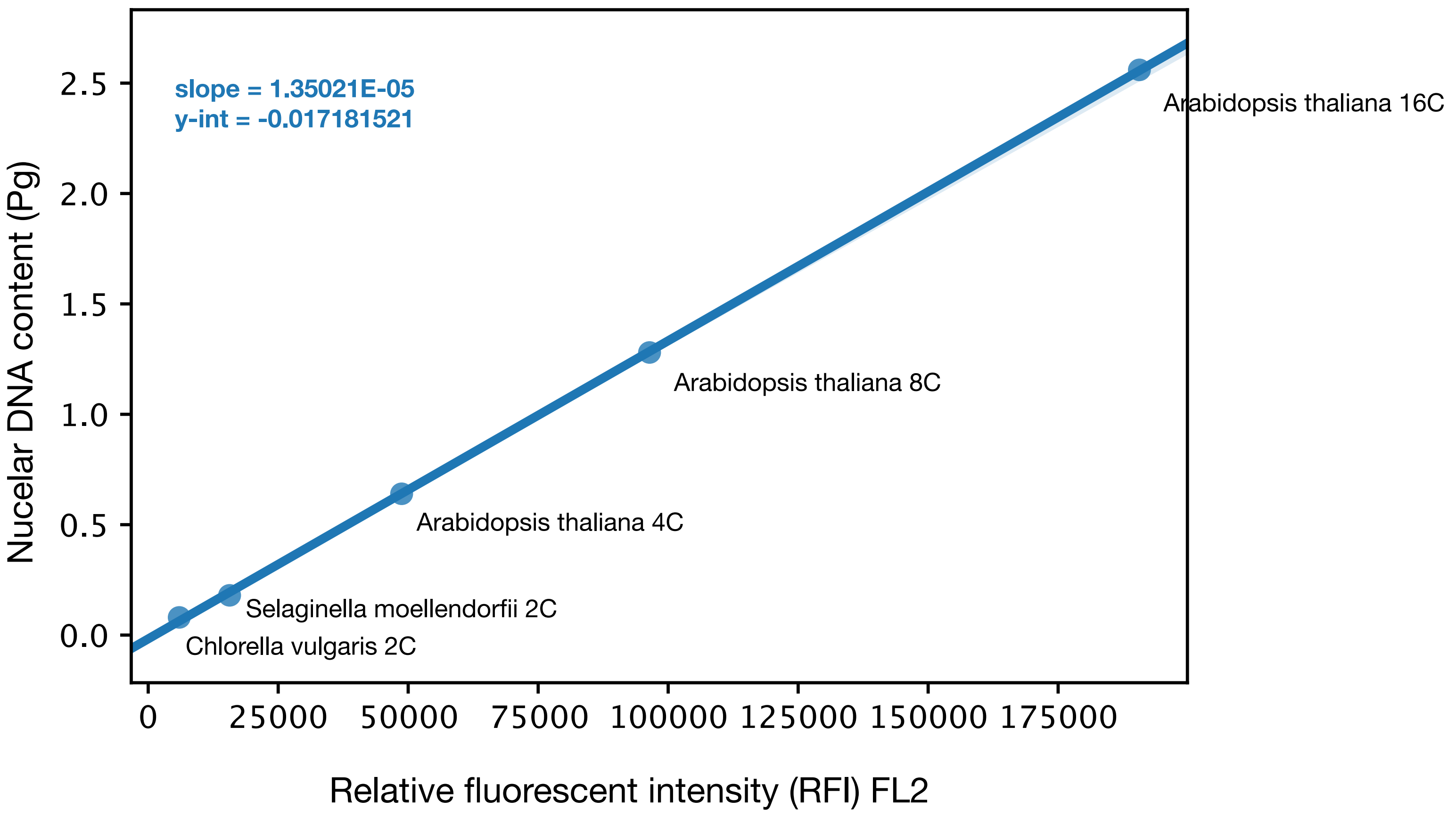


**Fig. S16. Estimation of nuclear DNA content.** Line of best fit through genome calibration standards with known genome sizes. The endoreduplicative nature of *A. thaliana* enabled identification of 4C, 8C, 16C, and in some cases 32C nuclei in this species (see SI ref 77). Relative fluorescent intensity of the FL2 channel measured RFI of propidium iodide-stained nuclei. Linear model was used to estimate the estimated nuclear DNA content in picograms for *Prymnesium parvum* strain.

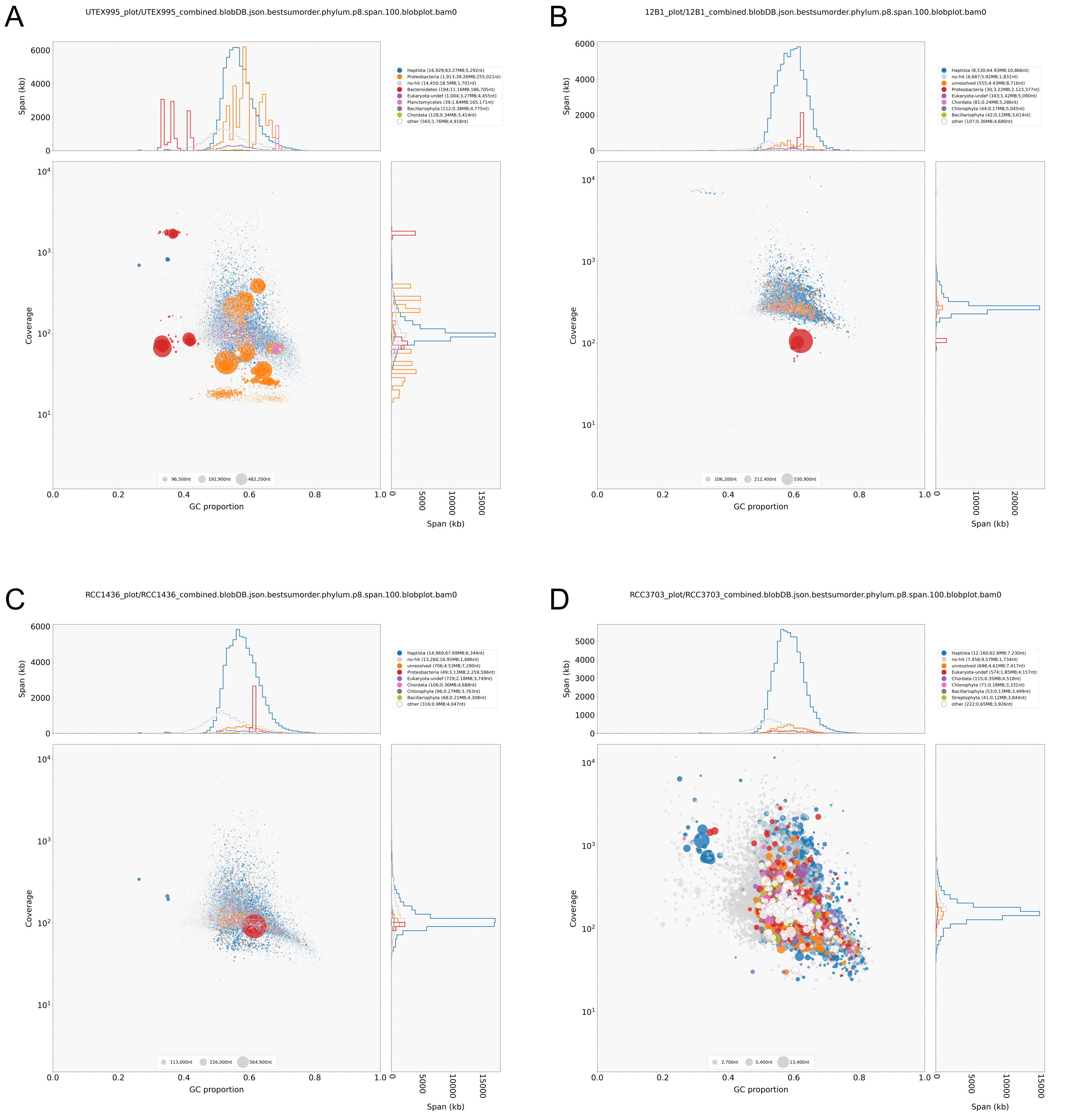


**Fig. S17. Example blobplots showing taxonomic breakdown of Abyss short-read assemblies.** For all blobplots, contigs with majority hits to haptophyte protein and/or nucleotide sequences are colored blue. A) UTEX 995 blobplot showing significant contamination from bacteriodetes (contigs depicted as red circles) and proteobacteria (orange circles). B) 12B1 and C) RCC1436 blobplots had some proteobacterial contamination as indicated by the red circles. D) The blobplot for RCC3703 indicated no bacterial contamination; all contigs had majority hits to eukaryotes, primarily haptophyte and other algal lineages. For all strains, contigs with majority hits to non-eukaryotes were excluded from further analysis. Reads that mapped to excluded contigs were also removed.

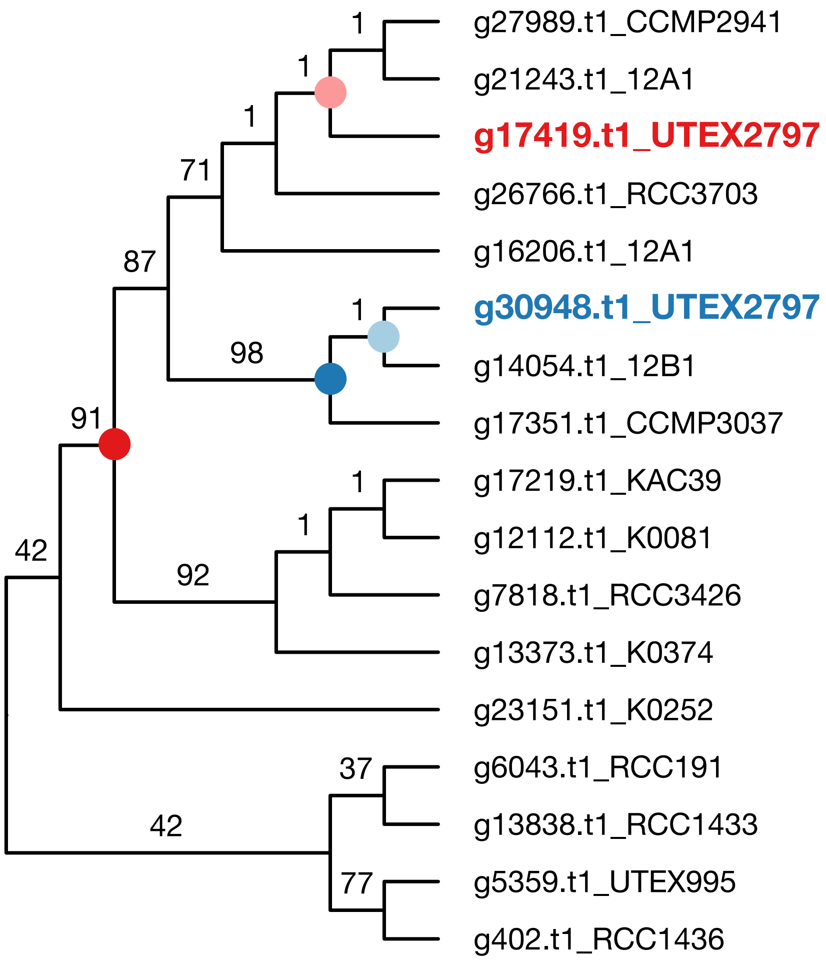


| Support threshold | Gene | Ancestral node | Ancestral node support | UTEX 2797 sister taxa | Designation |
|---|---|---|---|---|---|
| 0 | g30948.t1_UTEX2797 | | 1 | 12B1 | A1 gene |
| 0 | g17419.t1_UTEX2797 | | 1 | 12A1, CCMP2941 | neither |
| 90 | g30948.t1_UTEX2797 | | 98 | CCMP3037, 12B1 | A1 gene |
| 90 | g17419.t1_UTEX2797 | | 91 | K0081, RCC3426, CCMP2941, RCC3703, K0374, CCMP3037, KAC39, 12B1, 12A1 | neither |

**Fig. S18. Example Bio.Phylo tree parsing workflow.** For each UTEX 2797 gene in a gene tree, progress upwards toward the root until reaching an ancestral node that meets the ultrafast bootstrap support threshold; all descendants of that ancestral node, excluding UTEX 2797 are considered sister taxa to UTEX 2797. For the case of g17419 in UTEX 2797, when the support threshold is set to 0 (i.e., no threshold imposed), the ancestral node is designated by the pink circle, and the sister taxa are 12A1 and CCMP2941. When a support threshold >= 90 is imposed the node designated by the red circle is the ancestral node, and the sister taxa are K-0081, RCC3426, CCMP2941, RCC3703, K-0374, CCMP3037, KAC-39, 12B1, and 12A1. For the case of g30948, when the support threshold is set to 0 (i.e., no threshold imposed), the ancestral node is designated by the light blue circle, and the sister taxon is 12B1. When a support threshold >= 90 is imposed the node designated by the dark blue circle is the ancestral node, and the sister taxa are CCMP3037, and 12B1. UTEX 2797 genes whose sister taxa included 12B1 and/or CCMP3037 and no other strains were classified A1 genes. UTEX 2797 genes whose sister taxa included CCMP2941 and/or RCC3703 and no other strains were classified A2 genes.

**Table S1 (separate file).** Flow cytometry results.

**Table S2 (separate file).** Summary of *Prymnesium parvum* genome assemblies.

**Table S3 (separate file).** Summary of haptophyte genome assembly statistics.

**Table S4 (separate file).** Coordinates of each predicted telomeric repeat for 12B1 and UTEX 2797 assemblies.

**Table S5 (separate file).** Repeat content in 12B1 and UTEX 2797 assemblies.

**Table S6 (separate file).** Summary of gene model statistics for Prymnesium parvum genome assemblies.

**Table S7 (separate file).** Summary of BUSCO analysis performed on assemblies.

**Table S8 (separate file).** Strain information.

**Table S9 (separate file).** Status of Eukaryota BUSCO genes in haptophyte genome assemblies.

**Table S10 (separate file).** Median synonymous substitutions per synonymous site (Ks).

**Table S11 (separate file).** Breadth of coverage (BOC) analysis.

**Table S12 (separate file).** GRAMPA gene tree-Species tree reconcilation.

**Table S13 (separate file).** Phylogenetic analysis of UTEX 2797 haplotypes.

**Table S14 (separate file).** Summary of nuclear DNA content, haploid genome size, and cell volume.

**Table S15 (separate file).** OrthoFinder results.

**Table S16 (separate file).** Matrix of shared orthogroups between strains.

**Table S17 (separate file).** Tests for enrichment of GO and KEGG gene annotation terms.

**Table S18 (separate file).** Summary of phylogenetic screen for horizontal gene transfer.

**Table S19 (separate file).** Eleven HGTs with strong phylogenetic support.

**Table S20 (separate file).** Description of culture conditions for RNA-seq for gene calling.
